# Colon Cancer Cells Evade Drug Action by Enhancing Drug Metabolism

**DOI:** 10.1101/2023.12.21.572817

**Authors:** Bojie Cong, Teena Thakur, Alejandro Huerta Uribe, Evangelia Stamou, Sindhura Gopinath, Oliver Maddocks, Ross Cagan

## Abstract

Colorectal cancer (CRC) is the second most deadly cancer worldwide. One key reason is the failure of therapies that target RAS proteins, which represent approximately 40% of CRC cases. Despite the recent discovery of multiple alternative signalling pathways that contribute to resistance, durable therapies remain an unmet need. Here, we use liquid chromatography/ mass spectrometry (LC/MS) analyses on *Drosophila* CRC tumour models to identify multiple metabolites in the glucuronidation pathway—a toxin clearance pathway—as upregulated in trametinib-resistant *RAS/APC/P53* (“*RAP*”) tumours compared to trametinib-sensitive *RAS^G12V^* tumours. Elevating glucuronidation was sufficient to direct trametinib resistance in *RAS^G12V^*animals while, conversely, inhibiting different steps along the glucuronidation pathway strongly reversed *RAP* resistance to trametinib. For example, blocking an initial HDAC1-mediated deacetylation step with the FDA-approved drug vorinostat strongly suppressed trametinib resistance in *Drosophila RAP* tumours. We provide functional evidence that pairing oncogenic RAS with hyperactive WNT activity strongly elevates PI3K/AKT/GLUT signalling, which in turn directs elevated glucose and subsequent glucuronidation. Finally, we show that this mechanism of trametinib resistance is conserved in an *KRAS/APC/TP53* mouse CRC tumour organoid model. Our observations demonstrate a key mechanism by which oncogenic RAS/WNT activity promotes increased drug clearance in CRC. The majority of targeted therapies are glucuronidated, and our results provide a specific path towards abrogating this resistance in clinical trials.s

## Introduction

Despite recent advances in RAS pathway therapies, RAS-mutant colorectal cancer (CRC) has proven poorly sensitive to most targeted CRC therapies in the clinics^1^. This is somewhat surprising, as RAS pathway inhibitors have shown strong efficacy in *RAS*-mutant CRC pre-clinical studies. One important factor that has emerged is the role of genetic complexity: drug resistance typically increases in preclinical models that are more genetically complex^2,3^. We recently reported this phenomenon in Drosophila CRC models, both preclinically and in complex fly avatars as a part of a clinical trial: compared to oncogenic RAS alone, additionally targeting tumour suppressors APC and P53 (“RAP”) consistently led to emergent drug resistance^3,4^. However, the mechanisms that link genetic complexity to resistance across a broad spectrum of targeted therapies remains poorly understood.

A growing number of studies have shown that genetic and signalling complexity play key roles in drug response. For example, genomic mutations that lead to amplification or ‘rewiring’ of key signalling pathways have been linked to failure of targeted therapies^5,6^; however, co-targeting of these pathways has to date failed to yield durable KRAS-mutant CRC treatments^7,8,9,10^. An alternative possibility is emergence of a drug target-agnostic mechanism in response to genomic complexity. *KRAS*, *APC* and *TP53* are the most commonly mutated genes reported for human CRC^3,11^: adenoma progression is associated with loss of *APC* paired with oncogenic mutations in *KRAS*; malignant transformation is associated with additional mutations in *TP53*^12,13^. We therefore focused on this canonical multi-gene mutation profile commonly seen in human CRC tumours.

Oncogenic RAS is a key driver for tumour progression in up to 25% of all human cancers^14^. As a such, components of the RAS/MAPK pathway remain a high priority for targeted therapy. For example, Food and Drug Administration (FDA) approved MEK inhibitor trametinib proved effective in preclinical CRC models, but showed no therapeutic benefit in CRC patients^15,16^. Here, we report an LC/MS analysis comparing drug response in Drosophila *RAP* and *RAS^G12V^* hindgut tumours. We found that drug resistance in *RAP* tumours was primarily associated with upregulated drug metabolism via glucuronidation, a primary toxin clearance pathway used by cells to clear most cancer drugs. Blocking this upregulation had no direct effect on hindgut tumours, but restored drug sensitivity in *RAP* Drosophila and mouse organoid tumours to a level that mirrored tumours with *RAS^G12V^* alone. We further demonstrate that patient-accessible drugs such as vorinostat can block a key glucuronidation preparatory step to strongly sensitize tumours to trametinib. Together, our results demonstrate how a canonical CRC mutation profile elevates a key detoxification pathway to promote general drug resistance; interfering with this process provides a blueprint for sensitizing genetically complex tumours to targeted therapies.

## Results

### Glucuronidation pathway induces trametinib resistance in genetically complex tumours

The *Drosophila* hindgut has proven a useful tool for modelling CRC including for predicting therapeutics^4^. To identify the most effective inhibitor against *Ras^G12V^* tumours, we targeted transgenes to the developing hindgut using *byn-GAL4* and performed a limited FDA drug screen: the potent and specific MEK inhibitor trametinib was especially effective in reducing oncogenic *Ras^G12V^*-mediated transformation in the *Drosophila* hindgut, leading to increased animal survival (Figure S1a-b). Feeding larvae with 1 µM trametinib strongly rescued *byn>Ras^G12V^*-induced lethality (Figure 1a). In contrast, a multigenic *Ras^G12V^*, *Apc^RNAi^*, *P53^RNAi^* CRC model (*byn>RAP*)—designed to capture the three most common mutations reported for CRC—was resistant to trametinib both for animal survival (Figure 1a) and for transformation of the hindgut proliferative zone (HPZ). These data indicate an emergent resistance to trametinib in *byn>RAP* tumours, mirroring the trametinib resistance observed in *KRAS*-mutant CRC patients.

**Figure 1:**
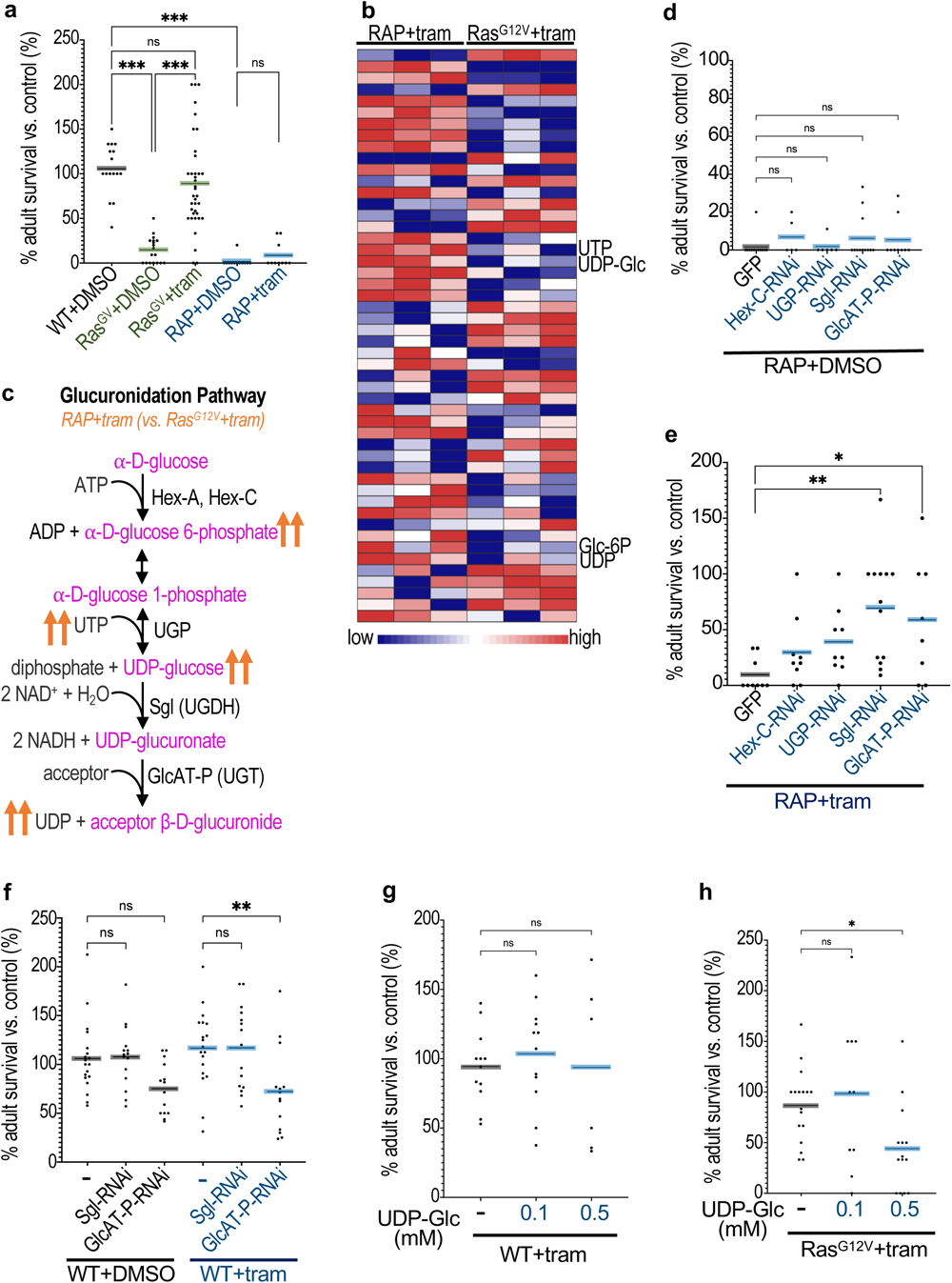
Glucuronidation pathway induces trametinib resistance in *Drosophila*. (A, D, E, F, G and H) Percent survival of transgenic flies to adulthood relative to control flies was quantified in the present or absence of trametinib (1μM) or UDP-Glc as indicated. (A) Wild-type (WT), *Ras^G12V^* and *RAP*; (D and E) *RAP + GFP* (control), *RAP +Hex-C-RNAi*, *RAP +UGP-RNAi*, *RAP +Sgl-RNAi*, *RAP + GlcAT-P-RNAi*; (F) *Sgl-RNAi* and *GlcAT-P-RNAi*; (G) WT; (H) *Ras^G12V^*. (B) A heatmap of LC/MS showed top 50 metabolises. (C) An overview of the glucuronidation pathway. Transgene expression was induced in *Drosophila* hindguts by a *byn-GAL4* driver. Drug concentrations indicate final food concentrations. Each data point represents a replicate.

Recent studies have linked metabolite changes to drug resistance in liver, lung, and renal cancer models^17,18,19^. To identify a metabolite fingerprint of emergent trametinib resistance, we performed a metabolomics analysis by liquid chromatography–mass spectrometry (LC– MS), comparing *byn>Ras^G12V^* and *byn>RAP* hindguts. 143 metabolites were altered in *byn>RAP* tumours upon administering trametinib (Figure 1b and Supplementary Table 1). An enrichment analysis (MetaboAnalyst 5.0, S1c) highlighted key differences between *byn>RAP* and *Ras^G12V^*response to trametinib, including transfer of acetyl groups into mitochondria (TAGIM), anaerobic glycolysis (Warburg effect), glutamate metabolism, citric acid cycle, nucleotide sugar metabolism, and purine metabolism (Figure S1c). The strongest enrichment was for metabolites associated with the glucuronidation pathway (Figure 1c), indicating upregulation of the pathway in *byn>RAP* tumours compared to *Ras^G12V^* tumours in the presence of trametinib. Upregulated metabolites included Glucose-6-phosphate (Glc-6P), UTP, UDP, and UDP-glucose (UDP-Glc; Figure 1b, 1c and S1d). Of note, glucuronidation is a key mechanism of drug resistance: cells use glucuronidation to solubilize and remove toxins including the majority of clinically relevant drugs^20^.

To investigate whether the glucuronidation pathway is essential for trametinib resistance in *byn>RAP* tumours, we used hindgut-targeted knockdown to reduce the activity of key glucuronidation enzymes including Hexokinase C (Hex-C; human ortholog: GCK), UDP-glucose Pyrophosphorylase (UGP; UGP2), Sugarless (Sgl; UDP-glucose 6-dehydrogenase, UDGH), and Glucuronyltransferase P (GlcAT-P; member of the human Glucuronosyltransferase family, UGT) (Figure 1c). Inhibiting the glucuronidation pathway promoted significant trametinib sensitivity in otherwise resistant *byn>RAP* tumours (Figure 1e). In particular, knockdown of Sgl or GlcAT-P significantly rescued tumour-induced lethality in the presence of trametinib (Figure 1e); neither knockdown impacted survival in the absence of trametinib (Figure 1d) or in wild type animals (Figure 1f). Conversely, elevating glucuronidation by supplementing the food with UDP-Glc—modelling elevated levels in RAP tumours—was sufficient to induce trametinib resistance in otherwise sensitive *byn>Ras^G12V^* tumours (Figure 1h). UDP-Glc did not affect survival of wild type animals (Figure 1g).

These data indicate that glucuronidation is both necessary and sufficient for emergent trametinib resistance in *byn>RAP* hindgut tumours. We next explored the mechanisms by which cancer gene combinations led to glucuronidation-dependent drug resistance.

### Pi3k/Akt signalling induces trametinib resistance by enhancing glucuronidation

Glucuronidation entails converting circulating glucose to intracellular UDP-glucuronide (UDP-Glc); transfer of glucuronide to a drug lead to its clearance coupled with UDP release (Figure 1c). *byn>RAP* hindguts displayed elevated glucose compared to *byn>Ras^G12V^* hindguts (Figure S1e), prompting us to investigate whether elevated glucose uptake led to increased glucuronidation. Our previous work showed that high dietary sugar (HDS) promoted glucose uptake in *Ras^G12V^ csk^-/-^* flies, enhancing tumour progression in eye-antennal epithelia as well as altering drug response^21,22,23^. Similarly, we found that HDS enhanced tumour progression in *byn>Ras^G12V^*hindguts resulting in increased animal lethality (Figure 2a); wild-type animals were not affected (Figure S2a). Importantly, HDS upregulated trametinib-dependent glucuronidation as determined by a UDP-glucose release assay (Figure 2b).

**Figure 2:**
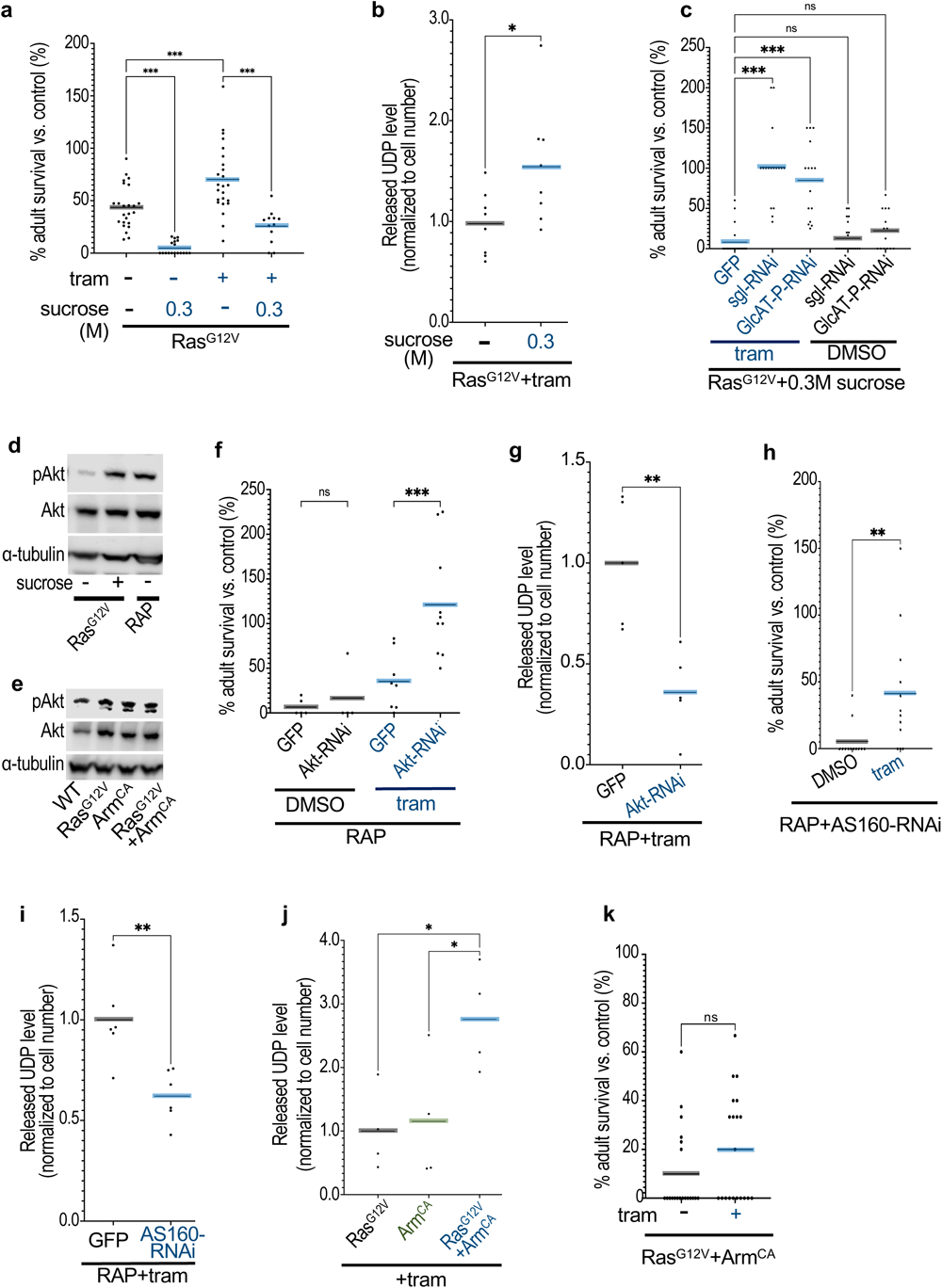
Pi3K/Akt signalling induces trametinib resistance by enhancing glucuronidation in *Drosophila*. (A, C, F, H and K) Percent survival of transgenic flies to adulthood relative to control flies was quantified in the present or absence of trametinib (1μM), sucrose. (A) *Ras^G12V^*; (C) *Ras^G12V^+GFP*, *Ras^G12V^+Sgl-RNAi* and *Ras^G12V^+GlcAT-P-RNAi*; (F) *RAP*+GFP, *RAP*+*Akt-RNAi*; (H) *RAP+AS160-RNAi*; (K) *Ras^G12V^*+*Arm^CA^* were induced in *Drosophila* hindguts. (D and E) Western blot analysis of *Drosophila* hindguts pAkt and Akt levels in *Ras^G12V^*, *RAP, WT, Arm^CA^*, *or Ras^G12V^*+*Arm^CA^*with or without sucrose. (G, I and J) Released UDP analysis of *RAP+GFP*, *RAP+Akt-RNAi*, *RAP+AS160-RNAi, Ras^G12V^*, *Arm^CA^*or *Ras^G12V^*+*Arm^CA^* with trametinib in *Drosophila* hindguts. Transgene expression was induced in *Drosophila* hindguts by a *byn-GAL4* driver. Increased dietary sugar led to increased glucuronidation and reduced trametinib activity, while targeting glucuronidation enzymes or Pi3K pathway activity strongly potentiated trametinib activity.

These results raised the question as to whether HDS directs drug resistance due to its impact on tumour progression *vs.* glucuronidation levels. Inhibiting glucuronidation by knockdown of key glucuronidation enzymes Sgl or GlcAT-P almost entirely suppressed the ability of HDS to reduce trametinib efficacy, but knockdown of either enzyme had no effect in the absence of trametinib (Figure 2c). Inhibiting glucuronidation did not impact wild-type animals even in the presence of HDS plus trametinib (Figure S2a). These data suggest that glucose uptake promotes trametinib resistance primarily by enhancing the glucuronidation pathway.

A key regulator of glucose uptake is Pi3k/Akt signalling, a Ras-initiated pathway that promotes glucose uptake by activating AS160 in mammalian cells^24^. Previous work indicates HDS promotes elevated glucose uptake by upregulation of phosphorylated Akt in normal *Drosophila*^25^. HDS also led to elevated Pi3k activity in *byn>Ras^G12V^* (*vs.* control) hindgut tumours as assessed by phosphorylated Akt (pAkt; Figure 2d). Compared to *byn>Ras^G12V^* alone, pAkt levels were strongly elevated in *byn>RAP* hindgut tumours even in the absence of HDS, phenocopying the effects of HDS (Figure 2d) and indicating that reducing Apc plus P53 further elevated Ras-dependent Pi3K activity. Knockdown of Akt or the AS160 ortholog *plx* strongly suppressed both glucuronidation and trametinib resistance in *byn>RAP* tumours (compare Figure 2f and h with Figures 1a, 2g and i). Further, activating Wnt pathway activity through the ß-catenin ortholog Arm was sufficient to strongly enhanced Pi3K activity and glucuronidation in the presence of Ras^G12V^, resulting in trametinib resistance in (normally sensitive) *byn>Ras^G12V^* tumours (Figure 2e, 2j; 2k compare with 1a). Wild-type animals were not affected (Figure S2b).

These data indicate that pairing elevated Ras plus Wnt pathway activities promotes trametinib resistance by (i) promoting glucose uptake in a Pi3K/Akt dependent manner, which in turn (ii) enhances glucuronidation and (iii) resistance to trametinib. Consistent with this view, pharmacological inhibition of Pi3K/Akt (LY294002) significantly increased trametinib sensitivity in *byn>RAP* animals (Figure S2c) at doses that did not impact wild-type animals (Figure S2d).

### Glucuronidation promotes trametinib resistance in mouse AKP organoids

To assess if our Drosophila data is relevant to mammalian CRC drug response, we investigated whether the mechanism of deacetylation plus glucuronidation promotes trametinib resistance in a murine colon cancer model. Using a mouse *VilCreER^T^*^2^, *Apc^fl/fl^*, *Kras^G12D/+^*, *Trp53^fl/fl^* (*AKP*) tumour organoid line derived from the small intestine, trametinib was strongly glucuronidated as assessed with a UDP-release assay (Figure 3a). *AKP* organoids were moderately sensitive to trametinib (Figure 3b-f). Consistent with our *Drosophila* results, promoting glucuronidation by adding UDP-Glc to the media inhibited response to high-dose trametinib (20 nM) in AKP tumour organoids (Figure S3a). Suppressing glucose uptake with (i) the GLUT1/GLUT4 inhibitor fasentin or (ii) the Pi3k/Akt inhibitors LY294002 and alpelisib significantly increased trametinib sensitivity. Single agents had no effect on tumour organoid expansion (Figure 3b-d, S3c, S3d, S3f and S3g).

**Figure 3:**
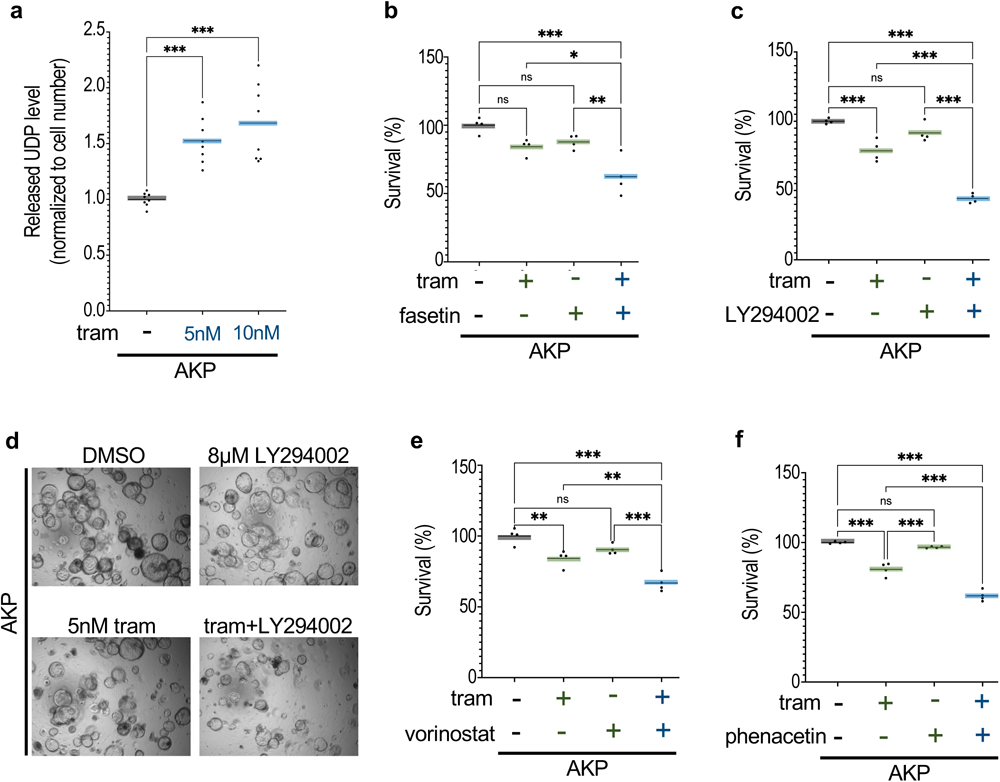
Deacetylation, glucuronidation lead to trametinib resistance in mouse AKP organoids. (A) Released UDP analysis of mouse AKP organoids in the present of trametinib. (B, C, E and F) Percent survival of AKP organoids relative to control was quantified in the present or absence of trametinib (5nM), fasentin (30μM), LY294002 (8μM), vorinostat (0.5μM) or phenacetin (100μM). (D) Representative images showing the impact of drugs on AKP organoids. Targeting glucuronidation led to increased effectiveness of trametinib.

### Trametinib glucuronidation was blocked by targeting deacetylation

Interfering with regulatory steps in the glucuronidation pathway including Pi3K signalling and glucuronidation enzymes potentiates trametinib activity in our CRC model. However, combining inhibition of these pathways with inhibition of MEK can lead to unwanted and significant toxicity^26,27,28,29,30,20^. In cancer patients, trametinib glucuronidation occurs in a two-step process: deacetylation followed by glucuronidation of the altered moiety^31^. Histone deacetylases (HDACs) deacetylate both histone proteins and non-histone cellular substrates that govern a wide array of disease processes including tumour progression and tumour therapy, and HDAC inhibitors are a staple of cancer treatment^32,33^. We therefore examined deacetylation as a potentially accessible therapeutic target.

We found that knockdown of the *Drosophila* deacetylase HDAC1 significantly suppressed trametinib glucuronidation in *byn>RAP* animals as determined by reduced UDP release (Figure 4a). The result was significantly increased sensitivity to trametinib and improved rescue of *byn>RAP* survival (Figure 4b). Similarly, co-feeding *byn>RAP* animals with the drug vorinostat (SAHA)—a clinically relevant inhibitor that binds to the active site of histone deacetylases^34^—significantly reduced trametinib resistance; vorinostat had no effect as a single agent (Figure 4c). This indicates that deacetylation is indeed required for glucuronidation and for trametinib resistance in *byn>RAP* flies. However, unlike glucuronidation, the baseline activity of HDAC did not differ between *byn>Ras^G12V^*and *byn>RAP* hindguts as assessed by a cell permeable, fluorescent HDAC substrate (Figure S4a). This suggests that, while deacetylation is necessary for glucuronidation, it likely does not account for the differential drug sensitivities observed between *byn>Ras^G12V^* and *byn>RAP* animals.

**Figure 4:**
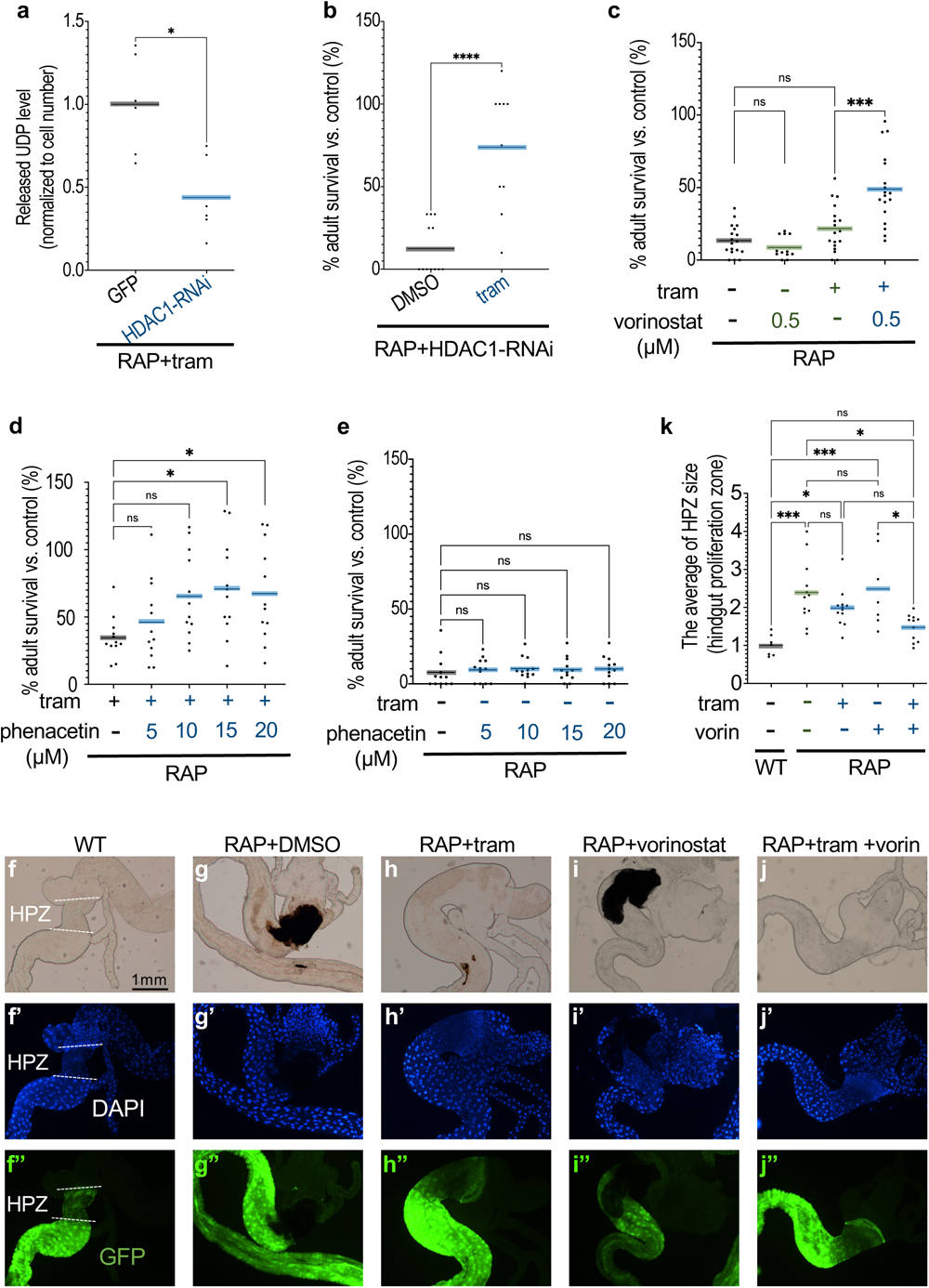
HDAC1 is required for glucuronidation of trametinib in *Drosophila*. (A) Released UDP analysis of *RAP+GFP* or *RAP+HDAC1-RNAi* with trametinib in *Drosophila* hindguts. (B, C, D and E) Percent survival of adult tumour flies relative to control flies was quantified in the present or absence of trametinib (1μM), vorinostat (0.5μM) or phenacetin. (B) *RAP +HDAC1-RNAi*, (C, D and E) *RAP*. Reduced HDAC activity led to reduced trametinib-dependent UDP release. (F-J) Images of the digestive tract of third instar larvae in the present or absence of trametinib (1 μM), vorinostat (0.5 μM) which include the hindgut proliferation zone (HPZ). Nuclei are visualized with 4′,6-diamidino-2-phenylindole (DAPI) staining, hindgut is marked by GFP. Scale bar 1mm. (K) The average of hindgut proliferation zone (HPZ) size was measured by Fiji ImageJ and quantified as relative size to wild-type (WT) hindgut. Transgene expression was induced in *Drosophila* hindguts by a *byn-GAL4* driver. Reducing deacetylation/glucuronidation with vorinostat increased trametinib’s ability to rescue hindgut size.

Similar to trametinib, the acetamide-based drug phenacetin is modified by deacetylation and glucuronidation^35^, providing a useful *in vivo* competitor for drug-modifying enzymes. Administered as a single agent, phenacetin had no affect on *byn>RAP* survival (Figure 4e). However, combining trametinib with phenacetin alleviated drug resistance to rescue animals in a dose-dependent manner (Figure 4d). This data further supports the view that, similar to human patients^31^, *byn>RAP* animals require a two-step modification to suppress trametinib: deacetylation followed by glucuronidation.

The impact of glucuronidation on drug response extended beyond animal survival. Targeting the hindgut proliferative zone (HPZ) in *byn>RAP* animals led to significant overgrowth compared to control animals (Figure 4g compared to 4f, quantified in 4k). Consistent with our adult survival assay, inhibiting deacetylation (vorinostat) in the presence of trametinib significantly suppressed overgrowth of the HPZ in *byn>RAP* tumours; trametinib or vorinostat alone did not have a strong effect (Figure 4h-j compared to 4g, quantified in 4k). Together, our data indicate that glucuronidation is enhanced by reducing Apc plus P53 activities in genotypically *byn>RAS^G12V^* hindguts, leading to emergent drug resistance.

Finally, inhibiting trametinib deacetylation by HDAC inhibitor (vorinostat) or via a competing substrate (phenacetin) significantly suppressed trametinib resistance in mouse AKP tumour organoids; again, single agents had no affect in the absence of trametinib (Figure 3e and 3f, S3e and S3h). These data indicate that, similar to fly RAP, deacetylation and glucuronidation are required for trametinib resistance in mouse AKP tumour organoids. HDAC inhibitors are well tolerated in the clinics, and this data provides a clinically accessible route to blocking glucuronidation of drugs such as trametinib that require a two-step modification.

## Discussion

Drug resistance in CRC patients remains one of the cancer field’s most persistent challenges. In this study, we demonstrate a mechanism by which CRC tumours achieve resistance to targeted therapies by elevating drug metabolism. We focused on trametinib, a potent MEK inhibitor that consistently failed to show significant clinical efficacy in KRAS-mutant CRC patients. We confirmed that trametinib is first deacetylated by Histone Deacetylase 1 (HDAC1) to prepare the drug for glucuronidation, which in turn resulted in inactivation/elimination of trametinib in a *byn>RAP* hindgut tumour model. This upregulation of glucuronidation was achieved by elevated RAS plus WNT pathway activities, which in turn increased glucose uptake in a Pi3K/AKT-dependent manner. Blocking this RAS-WNT-Pi3K-deacetylation/glucuronidation network at any one of several points strongly suppressed drug resistance in *byn>RAP* tumours (Figure 5). For example, combining trametinib with vorinostat proved potent in both Drosophila and mouse RAS-APC-P53 CRC models, addressing the most frequent three-mutation combination reported for CRC. Our findings indicate that glucuronidation—a major drug detoxification pathway—is upregulated in the context of oncogenic transformation and that this regulation is reversable.

**Figure 5:**
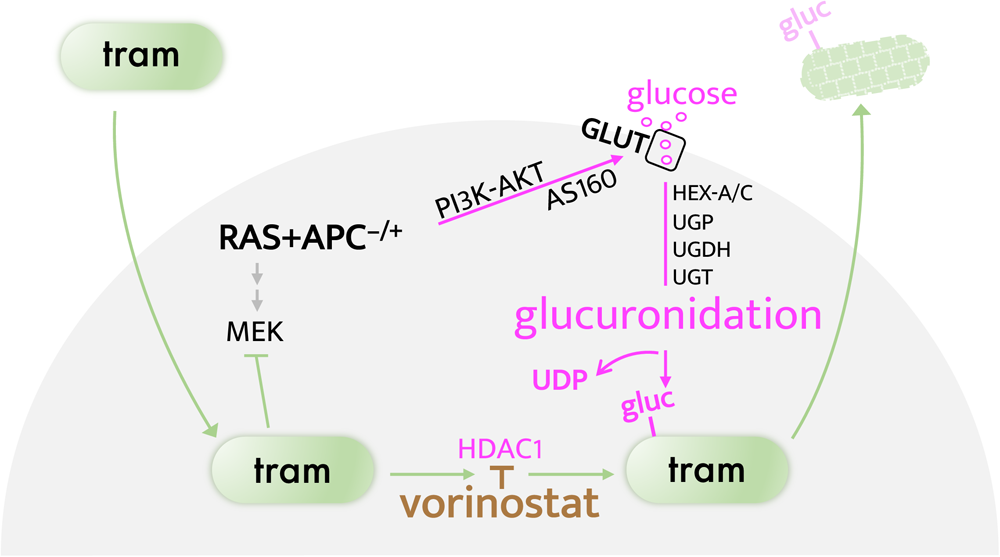
Schematic summary. Trametinib (tram) is a potent MEK inhibitor with the demonstrated preclinical ability to block RAS pathway signalling and oncogenic transformation. Pairing activated RAS and WNT activities leads to activation of PI3K/AKT signalling, AS160, and GLUT1/4 to increase glucose flux into cells. The result is elevated glucuronidation and elimination of trametinib. Potential therapeutic targets include HDAC1: deacetylation is an obligatory pre-step required for glucuronidation of some drugs including trametinib.

More than 70 therapeutic agents have been reported as metabolized by glucuronidation. Glucuronidation has been considered as a potential target of anticancer drug resistance including for colon cancer^36,20^, but mechanisms for regulating the pathway have been unclear and the large number of UDP-glucuronosyltransferases (UGTs) has made them poor candidates for targeting the pathway. Our study demonstrates that Ras/Erk plus Wnt/ß-catenin signalling upregulates the activity of Pi3K/Akt/Glut1, enhancing glucuronidation by promoting glucose uptake and promoting drug resistance in *byn>Ras^G12V^*tumours (Figure 5). Therapeutic targets include members of the Wnt/β-catenin and Pi3K/AKT pathways; for drugs such as trametinib that require an initial deacetylation step, we demonstrate the utility of HDAC inhibitors such as vorinostat as therapeutics. This two-step glucuronidation process also suggests a mechanism by which trametinib remains stable-but-inactive in the body: initial rapid deacetylation of trametinib keeps a metabolite in circulation until a slower glucuronidation step leads to its clearance. This view would explain previously-described distribution of metabolites in patients^31^.

Increased glucose uptake is a characteristic of cancer cells, and aerobic glycolysis efficiently produces ATP synthesis that promotes cell proliferation, known as the Warburg effect^37^. Glycolysis also mediates drug response including chemotherapeutics, immune checkpoint inhibitors and small molecule therapeutics through induction of autophagy, epithelial-mesenchymal transition (EMT), and by enhancing glycolytic enzymes impact on nonenzymatic activities^38,39^. Our data show that the high levels of glucose in transformed cells also activate glucuronidation, enhancing drug metabolism in canonical *RAS-APC-P53* CRC tumours. Of note, a high sugar diet was sufficient to activate glucuronidation in *Ras^G12V^* tumours, suggesting that high sugar diets can impact a patient’s response to anticancer drugs. In all, our study suggests multiple points to target along the emergent RAS-WNT-glucuronidation network for re-sensitizing tumours to targeted therapies, and provides insight into the long-observed difference between genetic and chemical deletion of a therapeutic target.

## Materials and methods

### *Drosophila* strains and genetics

Fly lines were cultured at room temperature or 25-29 °C on standard fly food or food-plus-compound. Fly food contained tayo agar 10g, soya flour 5g, sucrose 15g, glucose 33g, maize meal 15g, wheat germ 10g, treacle molasses 30g, yeast 35g, nipagin 10ml, propionic acid 5ml in 1000 ml water. Transgenes used (Bloomington Drosophila Stock Center number): *byn-gal4* (hindgut-specific line, V. Hartenstein), *UAS-Ras^G12V^* (second chromosome, G. Halder), *tub-gal80^TS^* (Bloomington Drosophila Stock Center #7017), *w^11^*^18^ (#3605), *UAS-mCD8-GFP* (#5137), *UAS-Hex-C-RNAi* (#57404), *UAS-UGP-RNAi* (#50902), *UAS-GlcAT-P-RNAi* (#67771), *UAS-sgl-RNAi* (#65348), *UAS-Akt-RNAi* (#82957), *UAS-plx-RNAi* (AS160, #66313), UAS-HDAC1-RNAi (#36800), and UAS-Arm^S10^ (#4782).

### Construction of the Drosophila RAP model

As previously described^4^, a pWalium expression vector was engineered with three Multiple Cloning Sites (MCS) downstream of UAS responsive elements. The RAP model was designed as a single plasmid construct incorporating the following: (i) oncogenic mutant Ras85D^G12V^ in the first multiple cloning site (MCS), (ii) 4 short 21 bp hairpins targeted to downregulate Apc plus 4 to downregulate P53 as a single 8-mer hairpin cluster into the third MCS with micro-RNA and intron derived spacers and loop sequences as previously described^4^. The resulting plasmid was then stably inserted into the 2^nd^ chromosome attP40 genome ‘landing site’. The sequence for the P53-Apc 8-mer:

> 1 actctgaata gggaattggg aattgagatc tgttctagac catattcagc ctttgagagt tggacgttca gttcaagtct atagttatat tcaagcatat
>
> 101 agacttgaac tgaacgtcca gcgaaatctg gcgagacatc gagtagtgcc accaaaagtt agccgcgttg tggaaaatcc ccatattcag cctttgagag
>
> 201 tcaacgtgga cgttcagttc aatagttata ttcaagcata ttgaactgaa cgtccacgtt ggcgaaatct ggcgagacat cggagggaaa tggagaacgc
>
> 301 aaaaatccca ttataatgga accatattca gcctttgaga gtccggatga acaaggcctt caatagttat attcaagcat attgaaggcc ttgttcatcc
>
> 401 gggcgaaatc tggcgagaca tcgatgtgct tgatcgtaac tccatccaaa ctcgatatta acccatattc agcctttgag agttcggtgg ttattgcttc
>
> 501 agcatagtta tattcaagca tatgctgaag caataaccac cgagcgaaat ctggcgagac atcgacaaat aatgttgcaa taaccagttg aaaccaatgg
>
> 601 aatccatatt cagcctttga gagtctcaaa gttgtgcaac tcttatagtt atattcaagc atataagagt tgcacaactt tgaggcgaaa tctggcgaga
>
> 701 catcgaacta acccgttcac ctgcgacaat ttttaatcta ttttccatat tcagcctttg agagtctgga cgaccagctt cgatgatagt tatattcaag
>
> 801 catatcatcg aagctggtcg tccaggcgaa atctggcgag acatcgagac cacgatcgaa agaggaaaaa cggaaaacga acgaaccata ttcagccttt
>
> 901 gagagtaaag atggacaaga agtacgatag ttatattcaa gcatatcgta cttcttgtcc atctttgcga aatctggcga gacatcggga ctagttttca
>
> 1001 ttatttatca gccagcacca acaacaccat attcagcctt tgagagtgca gctaaagatg gacaagaata gttatattca agcatattct tgtccatctt
>
> 1101 tagctgcgcg aaatctggcg agacatcgtt ggtactcgag atagtttgta tgaaatattt atatttttag cggccgcaag aa

### Chemicals

Drugs and compounds were as follows: trametinib (Selleckchem or biorbyt), UDP-glucose (ab120384, Abcam), sucrose (S0389, Sigma), LY294002 (Selleckchem), vorinostat (Selleckchem), phenacetin (Selleckchem), and alpelisib (Selleckchem). Drug and compound stocks were diluted in DMSO or water; drugs were then mixed into standard fly food with final DMSO concentration 0.1% to prevent toxicity.

### Liquid chromatography–mass spectrometry (LC–MS) in *Drosophila*

The third instar larvae were dissected in 1x PBS and hindguts (30) were lysed in 0.2 ml lysis solvent (methanol/acetonitrile/H2O (50:30:20)) on ice, homogenised, spun at 13,000 rpm at 4 °C for 15 minutes, transferred to fresh 1.5 ml tubes, and stored at −80 °C. Supernatants were then analysed by LC-MS by separating using hydrophilic interaction liquid chromatography with a SeQuant ZIC-pHILIC column (2.1 × 150 mm, 5 μm) (Merck). Analytes were detected with high-resolution, accurate-mass mass spectrometry using an Orbitrap Exactive in line with an Accela autosampler.

### Statistical analysis

Eggs were collected for 24 hours in drug-containing food at 18 °C to minimize transgene expression during embryogenesis to prevent embryonic effects or lethality. After 3 days, tubes were transferred to the appropriate temperature to induce transgene expression; the number of surviving *Drosophila* adults was quantified after 2 weeks. Statistical analysis was performed using Prims9. N.S P(>0.12), * P(0.033), ** P(0.002), *** P(0.001), and **** P(<0.0001). All statistical data were summarized in Supplementary Table2. All detailed genotypes were summarized in Supplementary Table3.

### Western blot analysis of *Drosophila* hindguts

Third instar larvae were dissected in 1x PBS and hindguts (30) were put into 2x Laemmli sample buffer (2.1% SDS, 26.3% glycerol, 65.8 mM Tris-HCl (pH 6.8), 0.01% bromophenol blue, 5% 2-mercaptoethanol, BIO-RAD). After homogenization at 100 °C for 5 minutes, samples were centrifuged at 13,000 rpm for 10 minutes, and lysate supernatants were used for western blot analysis. For signal detection, SuperSignal West Pico PLUS Chemiluminescent Substrate (Thermo Scientific) was used. Primary antibodies used were as follows: anti-Phospho-Akt (p-Akt, Ser473, D9E, Rabbit, #4060, Cell Signaling, 1:500 in 5%BSA/TBST (0.1% Tween20 in TBS)), anti-Akt (pan, C67E7, Rabbit, #4691, Cell Signaling, 1:1000 in 5%BSA/TBST), monoclonal anti-α-Tubulin (mouse, T5168-2ML, SIGMA, 1:5000 in 5%BSA/TBST). Secondary antibodies used: anti-mouse IgG, HRP-linked antibody (1:2000 in 5%BSA/TBST, #7076S, Cell Signaling), anti-rabbit IgG, and HRP-linked antibody (1:2000 in 5%BSA/TBST, #7074S, Cell Signaling).

### Endogenous UDP release assay, glucose assay, and HDAC activity assay

Third instar larvae were dissected in 1x PBS and hindguts (2-3) were assayed in 1x PBS. Cell number was measured by CellTiter-Fluor^TM^ Cell Viability Assay kit (Promega). Endogenous UDP release was measured by UDP-Glo^TM^ Glycosyltransferase Assay kit (Promega). HDAC activity was detected by HDAC Cell-Based Activity Assay kit (#600150, Cayman). Glucose levels were detected with the Glucose Assay Kit (ab169559, Abcam): 10 hindguts were homogenized on ice in cold Glucose Assay Buffer.

### Imaging of the digestive tract of third instar larvae

Third instar larvae were dissected in 1x PBS and fixed with 4% paraformaldehyde for 25 min at room temperature, then washed 15 min in PBT (0.1% Triton X in 1xPBS). Samples were mounted with DAPI-containing SlowFade Gold Antifade Reagent (#S36939, Molecular Probes). Fluorescence images were visualized on a Lecia TSC SPE confocal microscope.

### 3D Cell culture

Organoid cultures were previously derived from mouse small intestines: *AKP* (*VilCreER^T^*^2^ *Apc^fl/fl^, Kras^G12D/+^, Trp53^fl/fl^*)^40^. Isolated crypts were resuspended in Matrigel (BD Bioscience, 356231), plated in six-well plates, and overlaid with ENR growth medium comprising advanced DMEM/F12 (12634010) supplemented with penicillin–streptomycin (15070063), 10 mM HEPES (15630056), 2 mM glutamine (25030081), N2 (17502048), B27 (17504044) (all from Gibco, Life Technologies), 100 ng/ml Noggin (250-38), 500 ng/ml R-Spondin (315-32) and 50 ng/ml EGF (315-09) (all from PeproTech).

AKP organoids were maintained in OCM medium (2.5ml EGF, 1ml Noggin, 1.5ml N2/B27, 45ml Advanced Serum Medium (1X, 500ml Advanced DMEM/F12, 5ml L-glutamine, 5ml 1M Hepes, 5ml Pen/Strep) in Growth Factor Reduced Matrigel (#356231). Cell suspensions were generated from organoids by mechanical disruption and seeded in 96– well plates in OCM medium ± drugs. After 48 hours, cell viability was measured by CellTiter-Blue® Cell Viability Assay kit (Promega). Endogenous UDP release was detected by UDP-Glo^TM^ Glycosyltransferase Assay kit (Promega).

## Acknowledgements

We thank the Cagan Laboratory members, Andrew Campbell, and Justin Bower for important discussions. We also thank the Bloomington Drosophila Stock Center, and Owen Sansom for AKP organoids. This work was generously supported by grants from the NIH (R01CA258736) and a Royal Society Wolfson Fellowship.

## Compliance with ethical standards

The authors declare that they have no conflict of interest.

## Supporting information

**Supplementary Figure 1:**
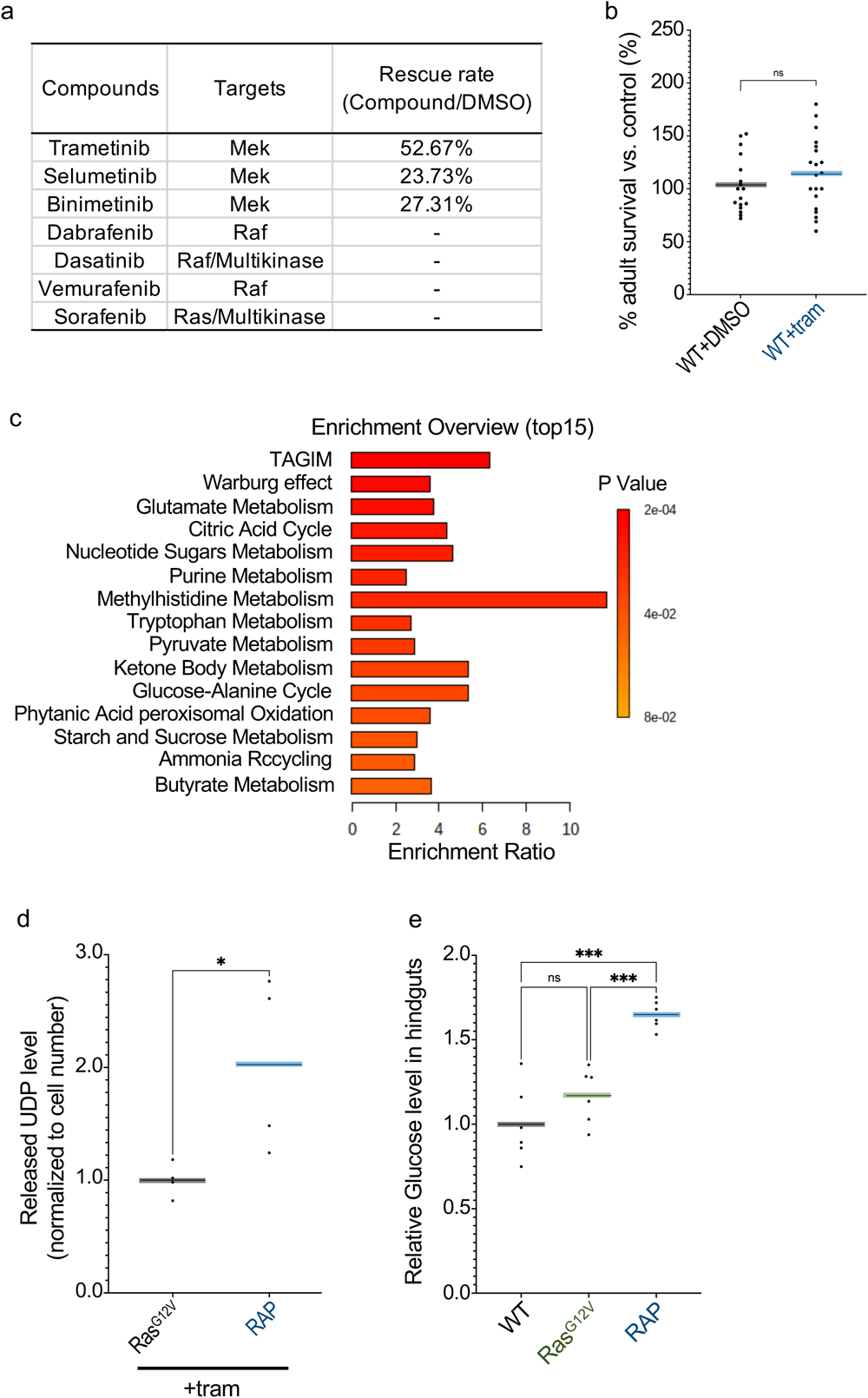
Screen for RAS pathway inhibitors. (a) A summary of rescue rate of RAS pathway inhibitors in *Ras^G12V^*hindgut tumours. (b) Percent survival of wild type flies to adulthood relative to control fly was quantified in the present or absence of trametinib (1 μM). (c) An enrichment overview of metabolites for *RAP* vs. *Ras^G12V^*in the presence of trametinib in fly hindguts. (d) Released UDP analysis of *Ras^G12V^* or *RAP* with trametinib in fly hindguts. (e) Relative glucose level of wild type (WT), *Ras^G12V^*, *RAP* in fly hindguts. Transgene expression was induced in *Drosophila* hindguts by a *byn-GAL4* driver.

**Supplementary Figure 2:**
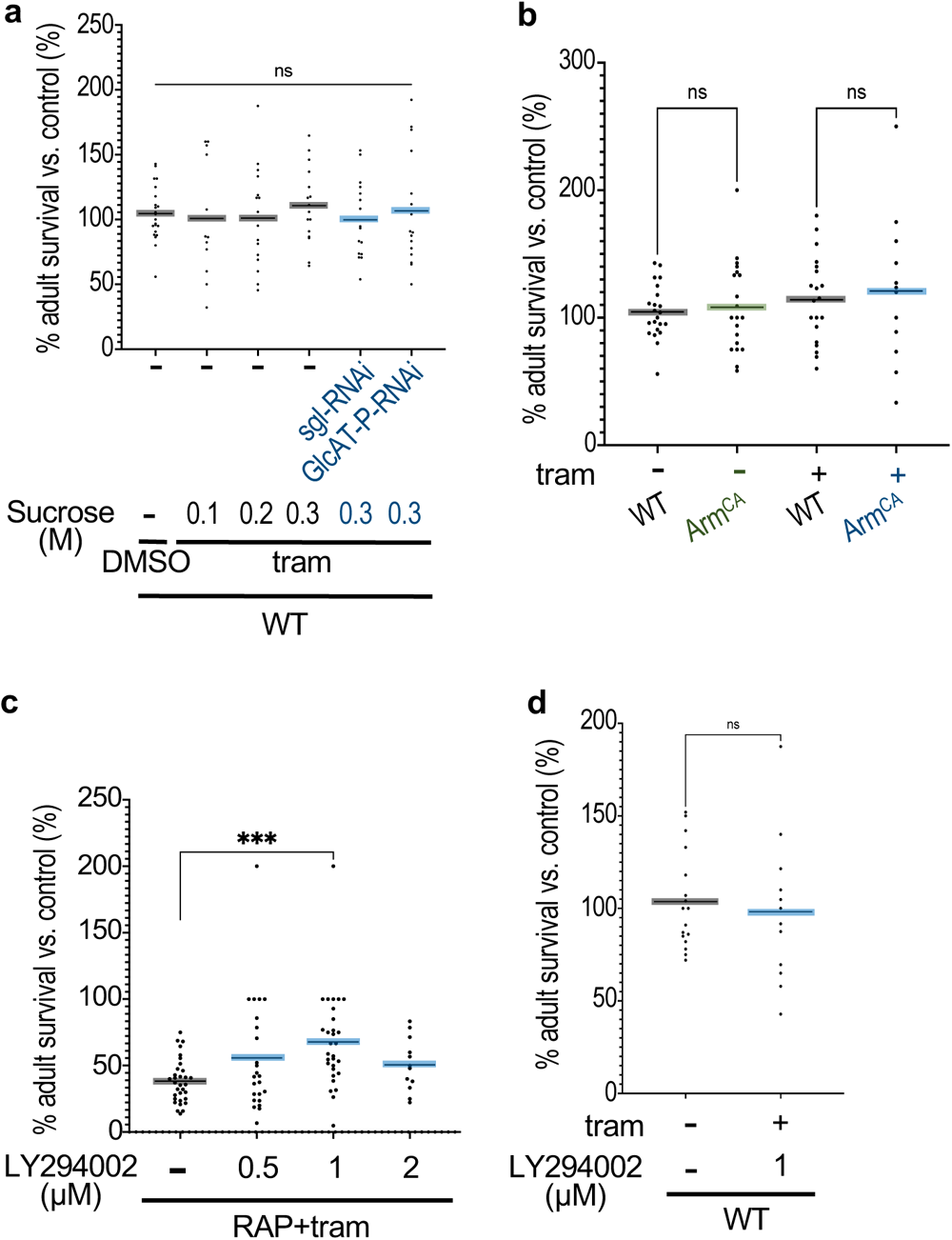
High dietary sucrose promoted glucuronidation. (a, b, c and d) Percent survival of transgenic CRC flies to adulthood relative to control fly was quantified in the present or absence of sucrose, trametinib (1μM), or LY294002. (a) wild-type (WT); (b) WT and *Arm^CA^*; (c) *RAP*; (d) WT.

**Supplementary Figure 3:**
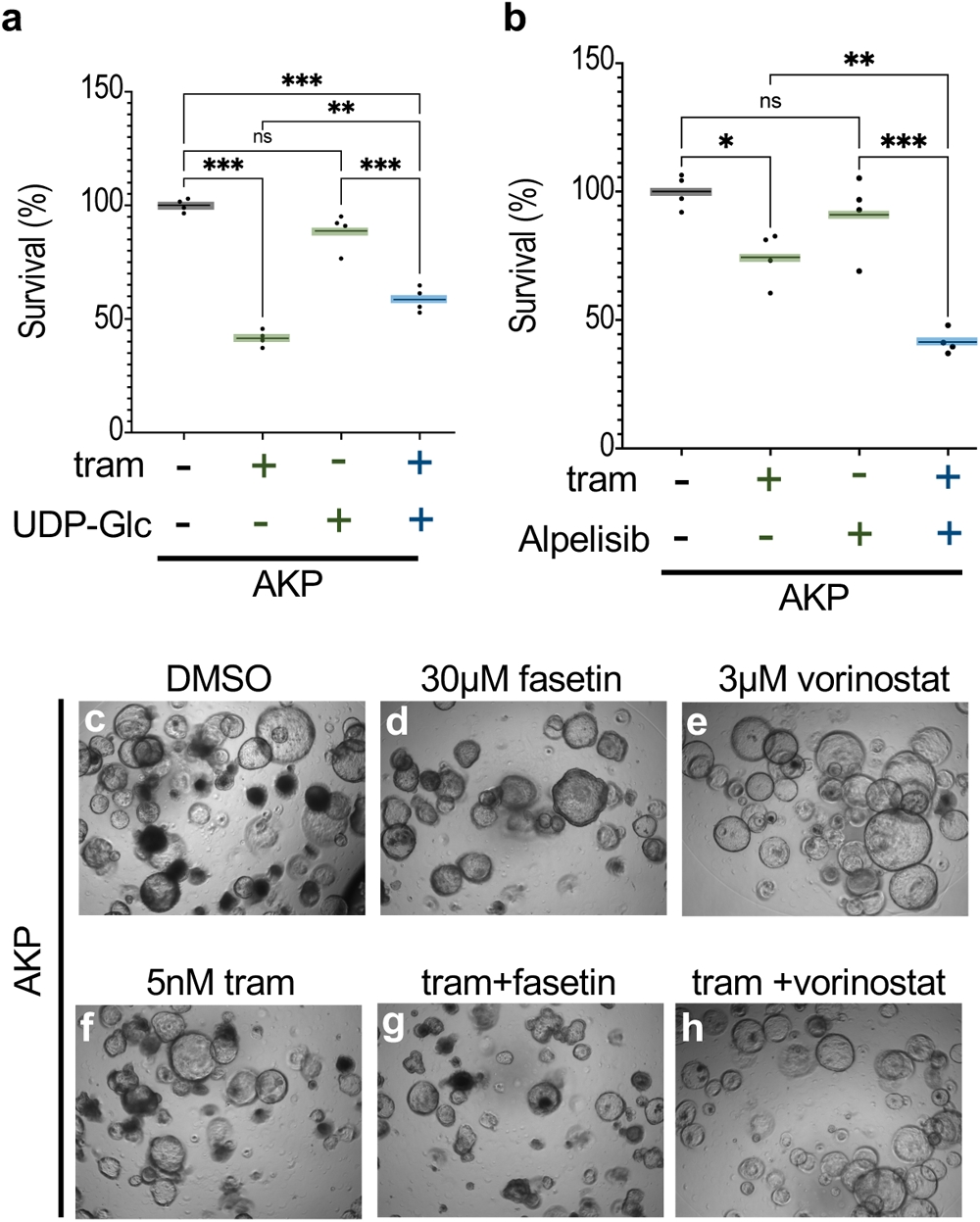
Deacetylation plus glucuronidation was required for trametinib resistance in mouse AKP organoids. (a and b) Percent survival of AKP organoids relative to control was quantified in the present or absence of trametinib (5nM), UDP-Glc (0.5μM), and alpelisib (2μM). (c-h) Representative images demonstrating the effect of drugs in AKP organoids.

**Supplementary Figure 4:**
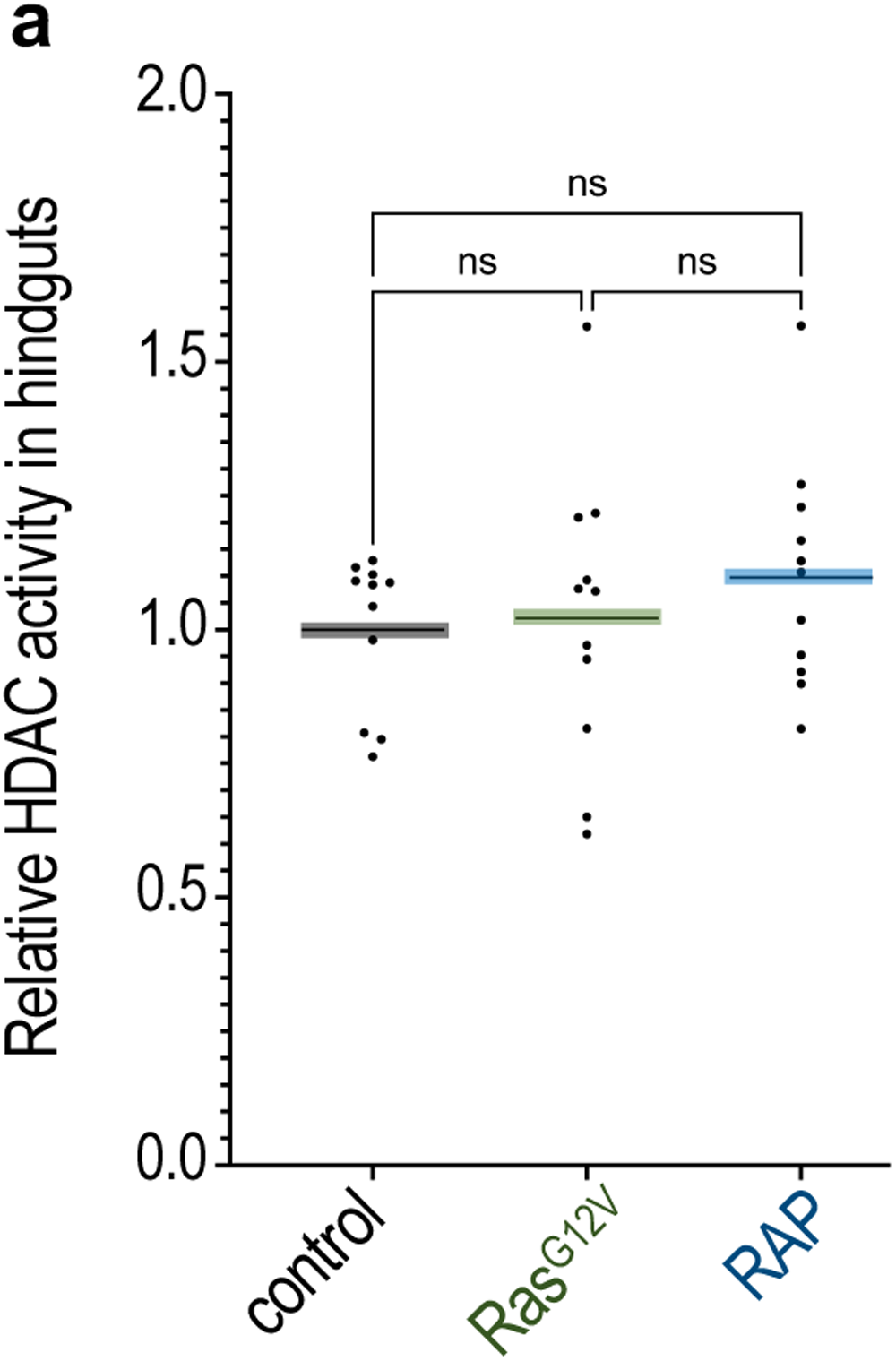
No difference observed in HDAC activity between *RAP* and *Ras^G12V^* in the fly hindgut. (a) HDAC activity analysis of wild type (WT), *Ras^G12V^* and *RAP* in fly hindguts.

**Supplementary Table 1:**
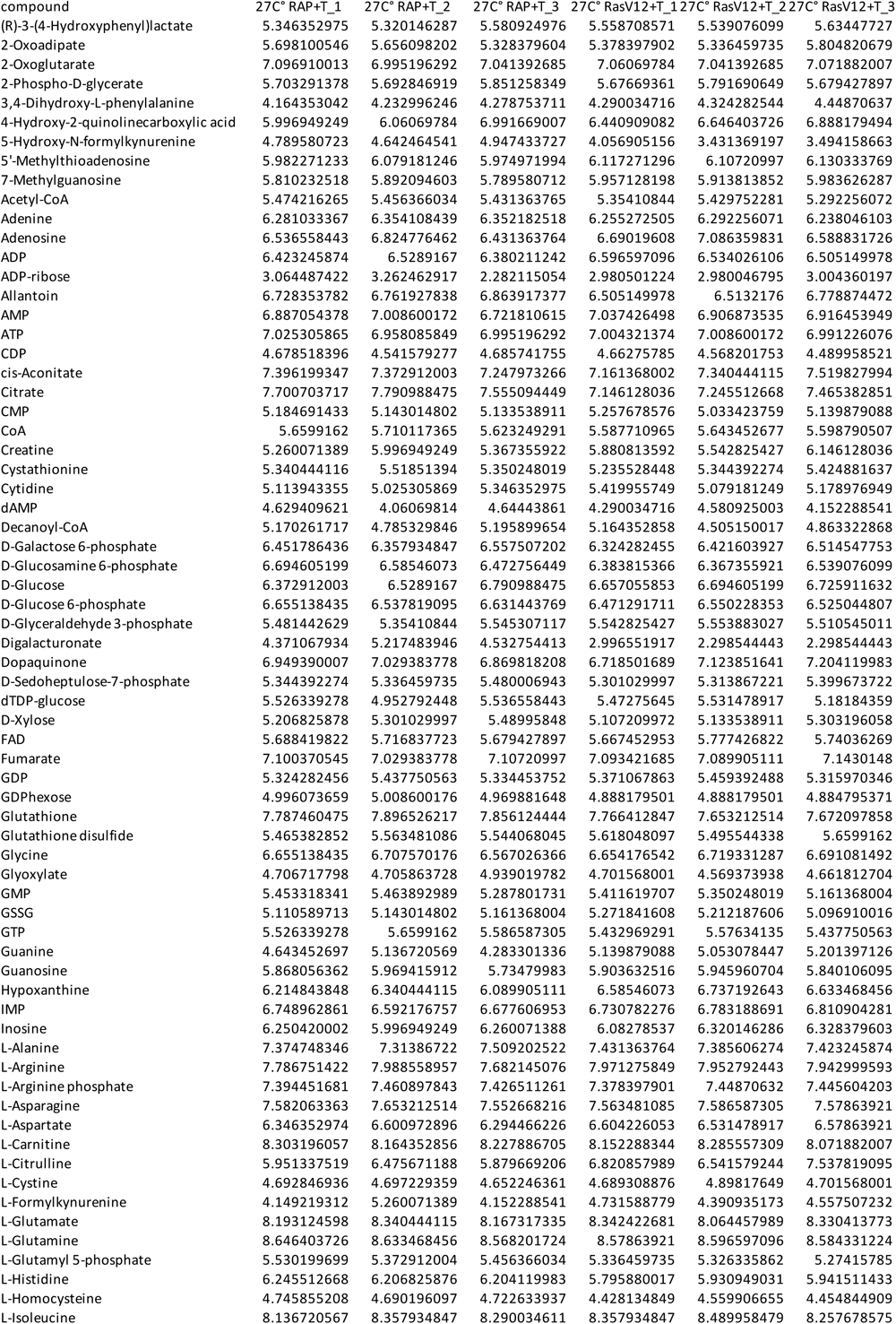
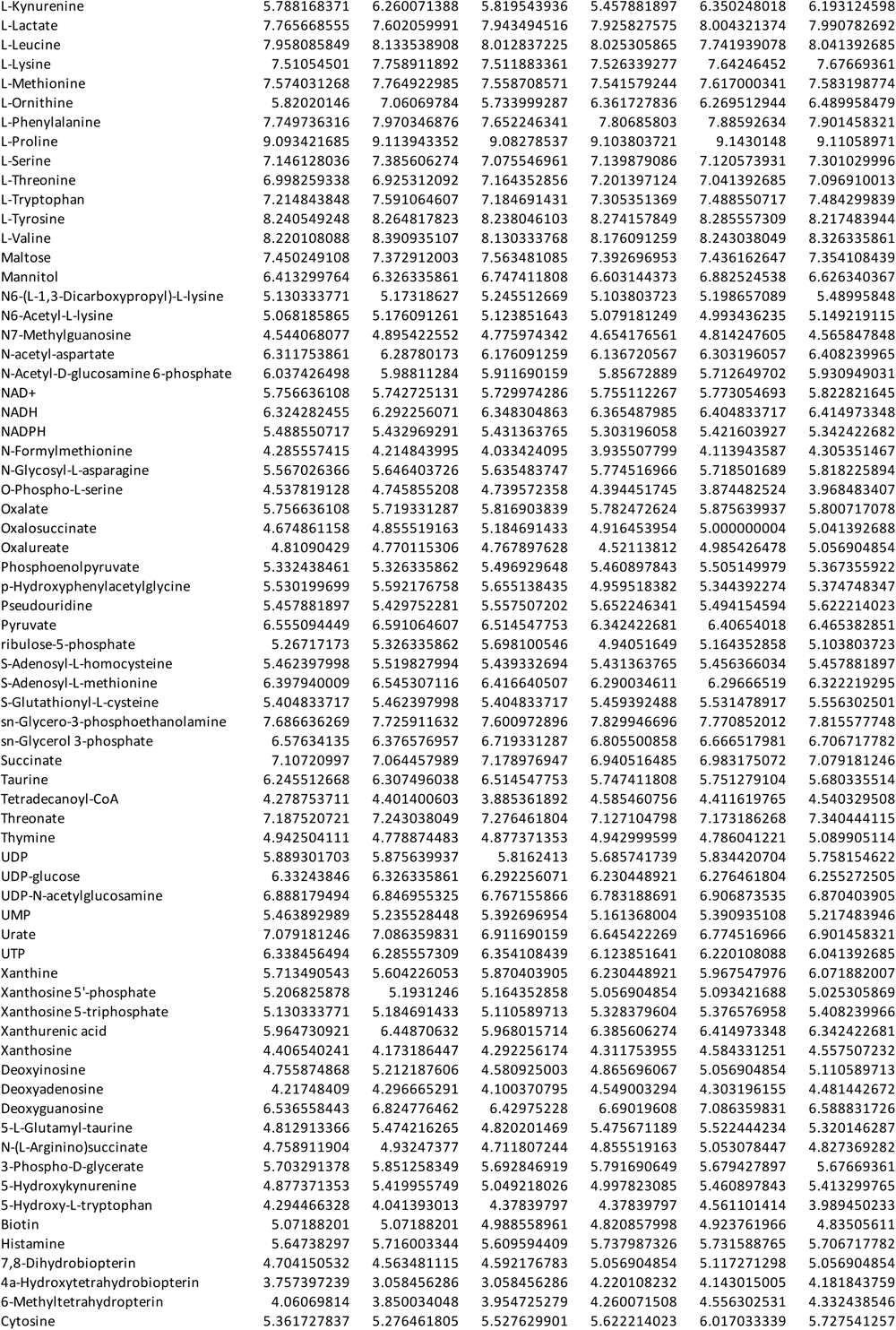
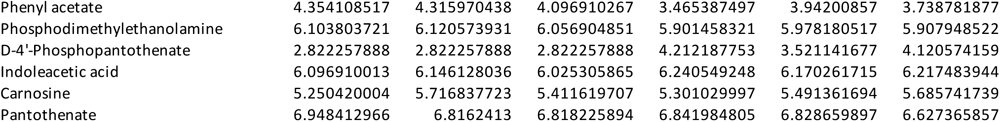
LC/MS data for *RAP* vs *Ras^G12V^* in the presence of trametinib in *Drosophila* hindguts.

**Supplementary Table 2:** Summary of statistical analyses.

| Figure1A | mean 1 | mean 2 | mean Diff | SE of Diff | n1 | n2 | Summary |
| --- | --- | --- | --- | --- | --- | --- | --- |
| WT_DMSO vs. RasG12V_DMSO | 105.6 | 14.71 | 90.91 | 12.62 | 16 | 18 | *** |
| WT_DMSO vs. RasG12V_T | 105.6 | 89.12 | 16.5 | 10.99 | 16 | 37 | ns |
| WT_DMSO vs. RAP_DMSO | 105.6 | 1.667 | 104 | 14.03 | 16 | 12 | *** |
| WT_DMSO vs. RAP_T | 105.6 | 8.666 | 96.96 | 14.81 | 16 | 10 | *** |
| RasG12V_DMSO vs. RasG12V_T | 14.71 | 89.12 | -74.41 | 10.56 | 18 | 37 | *** |
| RAP_DMSO vs. RAP_T | 1.667 | 8.666 | -6.999 | 15.73 | 12 | 10 | ns |
| Figure1D | mean 1 | mean 2 | mean Diff | SE of Diff | n1 | n2 | Summary |
| RAP_DMSO vs. +sgl-RNAi_DMSO | 1.667 | 6.25 | -4.583 | 3.777 | 12 | 12 | ns |
| RAP_DMSO vs. +μGP-RNAi_DMSO | 1.667 | 1.852 | -0.185 | 4.626 | 12 | 6 | ns |
| RAP_DMSO vs. +Hex-C-RNAi_DMSO | 1.667 | 6.858 | -5.191 | 4.924 | 12 | 5 | ns |
| RAP_DMSO vs. +GlcAT-P-RNAi_DMSO | 1.667 | 5.397 | -3.73 | 4.079 | 12 | 9 | ns |
| Figure1E | mean 1 | mean 2 | mean Diff | SE of Diff | n1 | n2 | Summary |
| RAP_Trmetinib vs. +sgl-RNAi_Trmetinib | 9.629 | 69.36 | -59.73 | 16.88 | 9 | 13 | ** |
| RAP_Trmetinib vs. +μGP-RNAi_Trmetinib | 9.629 | 39.06 | -29.43 | 18.35 | 9 | 9 | ns |
| RAP_Trmetinib vs. +Hex-C-RNAi_Trmetinib | 9.629 | 29.67 | -20.05 | 18.35 | 9 | 9 | ns |
| RAP_Trmetinib vs. +GlcAT-P-RNAi_Trmetinib | 9.629 | 58.75 | -49.12 | 18.92 | 9 | 8 | * |
| Figure1F | mean 1 | mean 2 | mean Diff | SE of Diff | n1 | n2 | Summary |
| WT_DMSO vs. sgl-RNAi_DMSO | 106.1 | 107.8 | -1.714 | 12.92 | 19 | 15 | ns |
| WT_DMSO vs. GlcAT-P-RNAi_DMSO | 106.1 | 75.17 | 30.93 | 13.18 | 19 | 14 | ns |
| WT_Trmetinib vs. sgl-RNAi_Trmetinib | 116.6 | 117 | -0.4274 | 12.92 | 19 | 15 | ns |
| WT_Trmetinib vs. GlcAT-P-RNAi_Trmetinib | 116.6 | 72.29 | 44.32 | 13.18 | 19 | 14 | ** |
| Figure1G | mean 1 | mean 2 | mean Diff | SE of Diff | n1 | n2 | Summary |
| WT_Trmetinib vs. +0.1mM μDP-G | 94.38 | 103.5 | -9.113 | 15.98 | 12 | 12 | ns |
| WT_Trmetinib vs. +0.5mM μDP-G | 94.38 | 93.65 | 0.7292 | 19.57 | 12 | 6 | ns |
| Figure1H | mean 1 | mean 2 | mean Diff | SE of Diff | n1 | n2 | Summary |
| RasG12V_Trmetinib vs. +0.1mM μDP-G | 86.67 | 98.57 | -11.91 | 19.78 | 17 | 10 | ns |
| RasG12V_Trmetinib vs. +0.5mM μDP-G | 86.67 | 44.16 | 42.51 | 17.91 | 17 | 14 | * |
| Figure2A | mean 1 | mean 2 | mean Diff | SE of Diff | n1 | n2 | Summary |
| RasG12V_DMSO vs. +0.3M Sp | 43.62 | 4.789 | 38.83 | 7.033 | 24 | 18 | *** |
| RasG12V_DMSO vs. RasV12_Trmetinib | 43.62 | 70.26 | -26.64 | 6.512 | 24 | 24 | *** |
| RasG12V_Trmetinib vs. +T+0.3M Sp | 70.26 | 25.62 | 44.64 | 7.975 | 24 | 12 | *** |
| Figure2B | median 1 | median 2 | Diff |  | n1 | n2 | Summary |
| RasG12V_Trmetinib vs. +0.3M Sp | 1 | 1.422 | 0.4222 |  | 8 | 8 | * |
| Figure2C | mean 1 | mean 2 | mean Diff | SE of Diff | n1 | n2 | Summary |
| RasG12V_Trmetinib+0.3M Sp vs. +sgl-RNAi_0.3M Sp | 8 | 12.86 | -4.857 | 10.19 | 20 | 21 | ns |
| RasG12V_Trmetinib+0.3M Sp vs. +sgl-RNAi_T+0.3M Sp | 8 | 101.4 | -93.37 | 10.75 | 20 | 17 | *** |
| RasG12V_Trmetinib+0.3M Sp vs. +GlcAT-P-RNAi_T+0.3M Sp | 8 | 84.78 | -76.78 | 11.14 | 20 | 15 | *** |
| RasG12V_Trmetinib+0.3M Sp vs. +GlcAT-P-RNAi_0.3M Sp | 8 | 22.32 | -14.32 | 11.36 | 20 | 14 | ns |
| Figure2F | mean 1 | mean 2 | mean Diff | SE of Diff | n1 | n2 | Summary |
| RAP+GFP_DMSO vs. RAP+akt-RNAi_DMSO | 6.666 | 18.08 | -11.42 | 23.64 | 5 | 8 | ns |
| RAP+GFP_Trmetinib vs. RAP+akt-RNAi_Trmetinib | 35.21 | 121.1 | -85.89 | 19.67 | 8 | 10 | *** |
| Figure2G | median 1 | median 2 | Diff |  | n1 | n2 | Summary |
| RAP+GFP_Trmetinib vs. RAP+akt-RNAi_Trmetinib | 1 | 0.3318 | -0.6682 |  | 5 | 5 | ** |
| Figure2H | median 1 | median 2 | mean Diff |  | n1 | n2 | Summary |
| RAP+AS160-RANI_DMSO vs. +Trmetinib | 0 | 26.79 | 26.79 |  | 12 | 12 | ** |
| Figure2I | median 1 | median 2 | Diff |  | n1 | n2 | Summary |
| RAP+GFP_Trmetinib vs. RAP+AS160-RNAi_Trmetinib | 0.9579 | 0.6207 | -0.3372 |  | 6 | 6 | ** |
| Figure2J | mean 1 | mean 2 | mean Diff | SE of Diff | n1 | n2 | Summary |
| RasG12V+ArmCA_Trmetinib vs. RasG12V_Trmetinib | 2.758 | 0.9995 | 1.758 | 0.586 | 4 | 4 | * |
| RasG12V+ArmCA_Trmetinib vs. ArmCA_Trmetinib | 2.758 | 1.154 | 1.604 | 0.586 | 4 | 4 | * |
| Figure2K | mean 1 | mean 2 | mean Diff | SE of Diff | n1 | n2 | Summary |
| RasG12V+ GFP_DMSO vs. + ArmCA_DMSO | 15 | 9.946 | 5.055 | 17.55 | 5 | 20 | ns |
| RasG12V+ GFP_DMSO vs. +GFP_Trmetinib | 15 | 82.31 | -67.31 | 16.99 | 5 | 29 | *** |
| RasG12V+ GFP_DMSO vs. +ArmCA_Trmetinib | 15 | 20 | -5 | 17.55 | 5 | 20 | ns |
| RasG12V+ ArmCA_DMSO vs. +ArmCA_Trmetinib | 9.946 | 20 | -10.05 | 11.1 | 20 | 20 | ns |
| RasG12V+GFP_Trmetinib vs. RasG12V+ArmCA_T | 82.31 | 20 | 62.31 | 10.2 | 29 | 20 | *** |
| Figure3A | median 1 | median 2 | Diff |  | n1 | n2 | Summary |
| RAP+GFP_Trmetinib vs. RAP+HDAC1-RNAi_Trmetinib | 1 | 0.3576 | -0.6424 |  | 6 | 6 | * |
| Figure3B | median 1 | median 2 | mean Diff |  | n1 | n2 | Summary |
| RAP+GFP_DMSO vs. RAP+HDAC1-RNAi_Trmetinib | 0 | 87.5 | 87.5 |  | 12 | 10 | **** |
| Figure3C | mean 1 | mean 2 | mean Diff | SE of Diff | n1 | n2 | Summary |
| RAP_DMSO vs. RAP_0.5μM Vorinostat | 13.47 | 8.677 | 4.788 | 6.018 | 18 | 12 | ns |
| RAP_DMSO vs. RAP_Trmetinib | 13.47 | 21.75 | -8.289 | 5.382 | 18 | 18 | ns |
| RAP_Trmetinib vs. RAP_T+ 0.5μM Vorinostat | 21.75 | 49 | -27.24 | 5.382 | 18 | 18 | *** |
| RAP_0.5μM Vorinosta vs. RAP_Trmetinib+ 0.5μM Vorinostat | 8.677 | 49 | -40.32 | 6.018 | 12 | 18 | *** |
| Figure3D | mean 1 | mean 2 | mean Diff | SE of Diff | n1 | n2 | Summary |
| RAP_Trmetinib vs. RAP_Trmetinib+5μM Phenacetin | 34.51 | 46.15 | -11.64 | 12.62 | 12 | 12 | ns |
| RAP_Trmetinib vs. RAP_Trmetinib+10μM Phenacetin | 34.51 | 65.4 | -30.89 | 12.62 | 12 | 12 | ns |
| RAP_Trmetinib vs. RAP_Trmetinib+15μM Phenacetin | 34.51 | 70.79 | -36.28 | 12.62 | 12 | 12 | * |
| RAP_Trmetinib vs. RAP_Trmetinib+20μM Phenacetin | 34.51 | 67.4 | -32.89 | 12.62 | 12 | 12 | * |

| Figure3E | mean 1 | mean 2 | mean Diff | SE of Diff | n1 | n2 | Summary |
| --- | --- | --- | --- | --- | --- | --- | --- |
| RAP_DMSO vs. RAP_5μM Phenacetin | 7.657 | 9.317 | -1.66 | 3.508 | 12 | 12 | ns |
| RAP_DMSO vs. RAP_10μM Phenacetin | 7.657 | 10.23 | -2.573 | 3.508 | 12 | 12 | ns |
| RAP_DMSO vs. RAP_15μM Phenacetin | 7.657 | 9.526 | -1.869 | 3.508 | 12 | 12 | ns |
| RAP_DMSO vs. RAP_20μM Phenacetin | 7.657 | 9.975 | -2.318 | 3.508 | 12 | 12 | ns |
| Figure3K | mean 1 | mean 2 | mean Diff | SE of Diff | n1 | n2 | Summary |
| WT vs. RAP_DMSO | 1 | 2.393 | -1.393 | 0.3056 | 7 | 12 | *** |
| WT vs. RAP_Tremetinib | 1 | 1.982 | -0.9819 | 0.3056 | 7 | 12 | * |
| WT vs. RAP_Vorinostat | 1 | 2.484 | -1.484 | 0.3435 | 7 | 7 | *** |
| WT vs. RAP_Tremetinib+Vorinostat | 1 | 1.475 | -0.4754 | 0.3107 | 7 | 11 | ns |
| RAP_DMSO vs. RAP_Tremetinib | 2.393 | 1.982 | 0.4112 | 0.2623 | 12 | 12 | ns |
| RAP_DMSO vs. RAP_Vorinostat | 2.393 | 2.484 | -0.09147 | 0.3056 | 12 | 7 | ns |
| RAP_DMSO vs. RAP_Tremetinib+Vorinostat | 2.393 | 1.475 | 0.9176 | 0.2682 | 12 | 11 | * |
| RAP_Tremetinib vs. RAP_Tremetinib+Vorinostat | 1.982 | 1.475 | 0.5064 | 0.2682 | 12 | 11 | ns |
| RAP_Vorinostat vs. RAP_Tremetinib+Vorinostat | 2.484 | 1.475 | 1.009 | 0.3107 | 7 | 11 | * |
| Figure4A | mean 1 | mean 2 | mean Diff | SE of Diff | n1 | n2 | Summary |
| AKP_DMSO vs. AKP_Tremetinib (5nM, 48h) | 1 | 1.525 | -0.5254 | 0.1169 | 8 | 8 | *** |
| AKP_DMSO vs. AKP_Tremetinib (10nM, 48h) | 1 | 1.684 | -0.6838 | 0.1169 | 8 | 8 | *** |
| Figure4B | mean 1 | mean 2 | mean Diff | SE of Diff | n1 | n2 | Summary |
| AKP_DMSO vs. AKP_Tremetinib (5nM, 48h) | 100 | 84.22 | 15.78 | 6.014 | 4 | 4 | ns |
| AKP_DMSO vs. AKP_Fasetin (30μM, 48h) | 100 | 87.82 | 12.18 | 6.014 | 4 | 4 | ns |
| AKP_DMSO vs. AKP_Tratinib+Fasetin | 100 | 62.4 | 37.61 | 6.014 | 4 | 4 | *** |
| AKP_Tremetinib (5nM, 48h) vs. AKP_Tremetinib+Fasetin | 84.22 | 62.4 | 21.82 | 6.014 | 4 | 4 | * |
| AKP_Fasetin (30μM, 48h) vs. AKP_Tremetinib+Fasetin | 87.82 | 62.4 | 25.42 | 6.014 | 4 | 4 | ** |
| Figure4C | mean 1 | mean 2 | mean Diff | SE of Diff | n1 | n2 | Summary |
| AKP_DMSO vs. AKP_Tremetinib (5nM, 48h) | 100 | 78.66 | 21.34 | 3.863 | 4 | 4 | *** |
| AKP_DMSO vs. AKP_LY294002 (8μM, 48h) | 100 | 91.59 | 8.41 | 3.863 | 4 | 4 | ns |
| AKP_DMSO vs. AKP_T+LY294002 (8μM, 48h) | 100 | 44.24 | 55.76 | 3.863 | 4 | 4 | *** |
| AKP_Tremetinib (5nM, 48h) vs. AKP_Tremetinib+LY294002 | 78.66 | 44.24 | 34.42 | 3.863 | 4 | 4 | *** |
| AKP_LY294002 (8μM, 48h) vs. AKP_Tremetinib+LY294002 | 91.59 | 44.24 | 47.35 | 3.863 | 4 | 4 | *** |
| Figure4E | mean 1 | mean 2 | mean Diff | SE of Diff | n1 | n2 | Summary |
| AKP_DMSO vs. AKP_Tremetinib (5nM, 48h) | 100 | 84.22 | 15.78 | 3.83 | 4 | 4 | ** |
| AKP_DMSO vs. AKP_Vorinostat (3μM, 48h) | 100 | 90.43 | 9.578 | 3.83 | 4 | 4 | ns |
| AKP_DMSO vs. AKP_Tremetinib+Vorinostat | 100 | 66.88 | 33.12 | 3.83 | 4 | 4 | *** |
| AKP_Tremetinib (5nM, 48h) vs. AKP_Tremetinib+Vorinostat | 84.22 | 66.88 | 17.34 | 3.83 | 4 | 4 | ** |
| AKP_Vorinostat (3μM, 48h) vs. AKP_Tremetinib+Vorinostat | 90.43 | 66.88 | 23.54 | 3.83 | 4 | 4 | *** |
| Figure4F | mean 1 | mean 2 | mean Diff | SE of Diff | n1 | n2 | Summary |
| AKP_DMSO vs. AKP_Tremetinib (5nM, 48h) | 100 | 80.82 | 19.18 | 2.191 | 4 | 4 | *** |
| AKP_DMSO vs. AKP_Phenacetin (100μM, 48h) | 100 | 96.61 | 3.393 | 2.191 | 4 | 4 | ns |
| AKP_DMSO vs. AKP_Tremetinib+Phenacetin | 100 | 61.84 | 38.16 | 2.191 | 4 | 4 | *** |
| AKP_Tremetinib (5nM, 48h) vs. AKP_Tremetinib+Phenacetin | 80.82 | 61.84 | 18.98 | 2.191 | 4 | 4 | *** |
| AKP_Phenacetin (100μM, 48h) vs. AKP_Tremetinib+Phenacetin | 96.61 | 61.84 | 34.77 | 2.191 | 4 | 4 | *** |
| Supplementary Figure1b | median 1 | median 2 | Diff |  | n1 | n2 | Summary |
| WT_DMSO vs. WT_Tremetinib | 100 | 114 | 14 |  | 17 | 20 | ns |
| Supplementary Figure1d | median 1 | median 2 | Diff |  | n1 | n2 | Summary |
| RasG12V_Tremetinib vs. RAP_Tremetinib | 1 | 2.047 | 1.047 |  | 4 | 4 | * |
| Supplementary Figure1e | mean 1 | mean 2 | mean Diff | SE of Diff | n1 | n2 | Summary |
| WT vs. RasG12V | 1 | 1.169 | -0.1691 | 0.09608 | 6 | 6 | ns |
| WT vs. RAP | 1 | 1.649 | -0.6489 | 0.09608 | 6 | 6 | *** |
| RasG12V vs. RAP | 1.169 | 1.649 | -0.4798 | 0.09608 | 6 | 6 | *** |
| Supplementary Figure2a | mean 1 | mean 2 | mean Diff | SE of Diff | n1 | n2 | Summary |
| WT_DMSO vs. WT_Tremetinib+0.1M Sucrose | 104.6 | 100.8 | 3.821 | 12.33 | 22 | 12 | ns |
| WT_DMSO vs. WT_Tremetinib+0.2M Sucrose | 104.6 | 101.1 | 3.517 | 11.29 | 22 | 16 | ns |
| WT_DMSO vs. WT_Tremetinib+0.3M Sucrose | 104.6 | 110.7 | -6.107 | 11.29 | 22 | 16 | ns |
| WT_DMSO vs. sgi-RNAi_Tremetinib+0.3M Sucrose | 104.6 | 99.77 | 4.851 | 11.29 | 22 | 16 | ns |
| WT_DMSO vs. GlcAT-P-RNAi_Tremetinib+0.3M Sucrose | 104.6 | 106.6 | -2.028 | 11.29 | 22 | 16 | ns |
| Supplementary Figure2b | mean 1 | mean 2 | mean Diff | SE of Diff | n1 | n2 | Summary |
| WT_DMSO vs. ArmCA_DMSO | 104.6 | 108.1 | -3.53 | 11.2 | 22 | 20 | ns |
| WT_Tremetinib vs. ArmCA_Tremetinib | 114.1 | 120.9 | -6.814 | 13.24 | 20 | 12 | ns |
| Supplementary Figure2c | mean 1 | mean 2 | mean Diff | SE of Diff | n1 | n2 | Summary |
| RAP_Tremetinib vs. RAP_Tremetinib+0.5μM LY294002 | 38.3 | 55.71 | -17.41 | 8.403 | 32 | 24 | ns |
| RAP_Tremetinib vs. RAP_Tremetinib+1μM LY294002 | 38.3 | 67.82 | -29.52 | 7.908 | 32 | 30 | *** |
| RAP_Tremetinib vs. RAP_Tremetinib+2μM LY294002 | 38.3 | 50.56 | -12.26 | 10.53 | 32 | 12 | ns |
| Supplementary Figure2d | median 1 | median 2 | Diff |  | n1 | n2 | Summary |
| WT_DMSO vs. WT_Tremetinib+LY29400 | 100 | 95.84 | -4.165 |  | 17 | 12 | ns |
| Supplementary Figure3a | mean 1 | mean 2 | mean Diff | SE of Diff | n1 | n2 | Summary |
| WT vs. RasG12V | 1 | 1.021 | -0.02124 | 0.09162 | 11 | 11 | ns |
| WT vs. RAP | 1 | 1.098 | -0.09776 | 0.09162 | 11 | 11 | ns |
| RasG12V vs. RAP | 1.021 | 1.098 | -0.07652 | 0.09162 | 11 | 11 | ns |
| Supplementary Figure4a | mean 1 | mean 2 | mean Diff | SE of Diff | n1 | n2 | Summary |
| AKP_DMSO vs. AKP_Tremetinib (20nM, 48h) | 100 | 41.55 | 58.45 | 3.842 | 4 | 4 | *** |
| AKP_DMSO vs. AKP_μDP-glc (160μM, 48h) | 100 | 88.72 | 11.28 | 3.842 | 4 | 4 | ns |
| AKP_DMSO vs. AKP_Tremetinib +μDP-glc | 100 | 58.57 | 41.44 | 3.842 | 4 | 4 | *** |
| AKP_Tremetinib (20nM, 48h) vs. AKP_Tremetinib +μDP-glc | 41.55 | 58.57 | -17.02 | 3.842 | 4 | 4 | ** |
| AKP_μDP-glc (160μM, 48h) vs. AKP_Tremetinib +μDP-glc | 88.72 | 58.57 | 30.16 | 3.842 | 4 | 4 | *** |
| Supplementary Figure4b | mean 1 | mean 2 | mean Diff | SE of Diff | n1 | n2 | Summary |
| AKP_DMSO vs. AKP_Trametinib (5nM, 48h) | 100 | 74.38 | 25.62 | 7.153 | 4 | 4 | * |
| AKP_DMSO vs. AKP_Alpelisib (2µM, 48h) | 100 | 90.99 | 9.015 | 7.153 | 4 | 4 | ns |
| AKP_DMSO vs. AKP_Trametinib (5nM) +Alpelisib (2µM, 48h) | 100 | 41.44 | 58.56 | 7.153 | 4 | 4 | *** |
| AKP_Trametinib (5nM, 48h) vs. AKP_Trametinib +Alpelisib | 74.38 | 41.44 | 32.94 | 7.153 | 4 | 4 | ** |
| AKP_Alpelisib (2µM, 48h) vs. AKP_Trametinib +Alpelisib | 90.99 | 41.44 | 49.54 | 7.153 | 4 | 4 | *** |

**Supplementary Table 3:**

Detailed genotypes.

**Figure 1:**

+/+ or Y; +/+; byn-Gal4, UAS-GFP, tub-Gal80^TS^/+ (a, f and g), +/+ or Y; UAS-Ras^G12V^/+; byn-Gal4, UAS-GFP, tub-Gal80^TS^/+ (a, b and h), +/+ or Y; UAS-Ras^G12V^, Apc-RNAi, UAS-P53-RNAi/+; byn-Gal4, UAS-GFP, tub-Gal80^TS^/+ (a and b), +/+ or Y; UAS-Ras^G12V^, Apc-RNAi, UAS-P53-RNAi/UAS-GFP; byn-Gal4, UAS-GFP, tub-Gal80^TS^/+ (d and e), +/+ or Y; UAS-Ras^G12V^, Apc-RNAi, UAS-P53-RNAi/UAS-Hex-C-RNAi; byn-Gal4, UAS-GFP, tub-Gal80^TS^/+ (d and e), +/+ or Y; UAS-Ras^G12V^, Apc-RNAi, UAS-P53-RNAi/UAS-UGP-RNAi; byn-Gal4, UAS-GFP, tub-Gal80^TS^/+ (d and e), +/+ or Y; UAS-Ras^G12V^, Apc-RNAi, UAS-P53-RNAi/UAS-Sgl-RNAi; byn-Gal4, UAS-GFP, tub-Gal80^TS^/+ (d and e), +/+ or Y; UAS-Ras^G12V^, Apc-RNAi, UAS-P53-RNAi/UAS-GlcAT-P-RNAi; byn-Gal4, UAS-GFP, tub-Gal80^TS^/+ (d and e), +/+ or Y; +/UAS-Sgl-RNAi; byn-Gal4, UAS-GFP, tub-Gal80^TS^/+ (f), +/+ or Y; +/UAS-GlcAT-P-RNAi; byn-Gal4, UAS-GFP, tub-Gal80^TS^/+ (f).

**Figure 2:**

+/+ or Y; UAS-Ras^G12V^/+; byn-Gal4, UAS-GFP, tub-Gal80^TS^/+ (a, b and d), +/+ or Y; UAS-Ras^G12V^/UAS-GFP; byn-Gal4, UAS-GFP, tub-Gal80^TS^/+ (c, e, j and k), +/+ or Y; UAS-Ras^G12V^/UAS-Sgl-RNAi; byn-Gal4, UAS-GFP, tub-Gal80^TS^/+ (c), +/+ or Y; UAS-Ras^G12V^/UAS-GlcAT-P-RNAi; byn-Gal4, UAS-GFP, tub-Gal80^TS^/+ (c), +/+ or Y; +/UAS-GFP; byn-Gal4, UAS-GFP, tub-Gal80^TS^/+ (c and e), +/+ or Y; UAS-Ras^G12V^, Apc-RNAi, UAS-P53-RNAi/+; byn-Gal4, UAS-GFP, tub-Gal80^TS^/+ (d), +/+ or Y; UAS-Ras^G12V^, Apc-RNAi, UAS-P53-RNAi/UAS-GFP; byn-Gal4, UAS-GFP, tub-Gal80^TS^/+ (f, g and i), +/+ or Y; UAS-Ras^G12V^, Apc-RNAi, UAS-P53-RNAi/UAS-Akt-RNAi; byn-Gal4, UAS-GFP, tub-Gal80^TS^/+ (f and g), +/+ or Y; UAS-Ras^G12V^, Apc-RNAi, UAS-P53-RNAi/UAS-AS160-RNAi; byn-Gal4, UAS-GFP, tub-Gal80^TS^/+ (h and i), UAS-Arm^CA^/+ or Y; UAS-GFP/+; byn-Gal4, UAS-GFP, tub-Gal80^TS^/+ (e, j and k), UAS-Arm^CA^/+ or Y; UAS-Ras^G12V^/+; byn-Gal4, UAS-GFP, tub-Gal80^TS^/+ (e, j and k).

**Figure 3:**

*VilCreERT2* Apc^fl/fl^, Kras^G12D/+^, Trp53^fl/fl^ (A-F).

**Figure 4:**

+/+ or Y; UAS-Ras^G12V^, Apc-RNAi, UAS-P53-RNAi/UAS-GFP; byn-Gal4, UAS-GFP, tub-Gal80^TS^/+ (a), +/+ or Y; UAS-Ras^G12V^, Apc-RNAi, UAS-P53-RNAi/UAS-HDAC1-RNAi; byn-Gal4, UAS-GFP, tub-Gal80^TS^/+ (a and b), +/+ or Y; UAS-Ras^G12V^, Apc-RNAi, UAS-P53-RNAi/+; byn-Gal4, UAS-GFP, tub-Gal80^TS^/+ (c, d, e and g-k), +/+ or Y; +/+; byn-Gal4, UAS-GFP, tub-Gal80^TS^/+ (f and k).

**Supplementary Figure 1:**

+/+ or Y; UAS-Ras^G12V^/+; byn-Gal4, UAS-GFP, tub-Gal80^TS^/+ (a, c, d and e), +/+ or Y; +/+; byn-Gal4, UAS-GFP, tub-Gal80^TS^/+ (b and e), +/+ or Y; UAS-Ras^G12V^, Apc-RNAi, UAS-P53-RNAi/+; byn-Gal4, UAS-GFP, tub-Gal80^TS^/+ (c, d and e).

**Supplementary Figure 2:**

+/+ or Y; +/+; byn-Gal4, UAS-GFP, tub-Gal80^TS^/+ (a, b and d), +/+ or Y; UAS-Ras^G12V^, Apc-RNAi, UAS-P53-RNAi/+; byn-Gal4, UAS-GFP, tub-Gal80^TS^/+ (c).

**Supplementary Figure 3:**

+/+ or Y; UAS-GFP/+; byn-Gal4, UAS-GFP, tub-Gal80^TS^/+ (a), +/+ or Y; UAS-Ras^G12V^/+; byn-Gal4, UAS-GFP, tub-Gal80^TS^/+ (a), +/+ or Y; UAS-Ras^G12V^, Apc-RNAi, UAS-P53-RNAi/+; byn-Gal4, UAS-GFP, tub-Gal80^TS^/+ (a).

**Supplementary Figure 4:**

*VilCreER^T^*^2^ *Apc*^fl/fl^, *Kras^G12D/+^*, *Trp53^fl/fl^* (a-h).

## References

1. Cagir, A. & Azmi, A. S. KRAS G12C inhibitors on the horizon. *Futur*. Med Chem. 11, 923–925 (2019).

2. Caponigro, G. & Sellers, W. R. Advances in the preclinical testing of cancer therapeutic hypotheses. Nat. Rev. Drug Discov. 10, 179–187 (2011).

3. Bangi, E., Murgia, C., Teague, A. G. S., Sansom, O. J. & Cagan, R. L. Functional exploration of colorectal cancer genomes using Drosophila. Nat. Commun. 7, 13615 (2016).

4. Bangi, E. et al. A personalized platform identifies trametinib plus zoledronate for a patient with KRAS-mutant metastatic colorectal cancer. Sci. Adv. 5, eaav6528 (2019).

5. Holohan, C., Schaeybroeck, S. Van, Longley, D. B. & Johnston, P. G. Cancer drug resistance: An evolving paradigm. Nat. Rev. Cancer 13, 714–726 (2013).

6. Kukal, S. et al. Multidrug efflux transporter ABCG2: expression and regulation. Cell. Mol. Life Sci. 78, 6887–6939 (2021).

7. Hallin, J. et al. The KRASG12C inhibitor MRTX849 provides insight toward therapeutic susceptibility of KRAS-mutant cancers in mouse models and patients. Cancer Discov. 10, 54–71 (2020).

8. Canon, J. et al. The clinical KRAS(G12C) inhibitor AMG 510 drives anti-tumour immunity. Nature 575, 217–223 (2019).

9. Biller, L. H. & Schrag, D. Diagnosis and treatment of metastatic colorectal cancer: A review. JAMA - J. Am. Med. Assoc. 325, 669–685 (2021).

10. Coupez, D., Hulo, P., Touchefeu, Y., Denis, M. G. & Bennouna, J. KRAS mutations in metastatic colorectal cancer: from a de facto ban on anti-EGFR treatment in the past to a potential biomarker for precision medicine. Expert Opin. Biol. Ther. 21, 1325–1334 (2021).

11. An, Y. et al. Clinicopathological and Molecular Characteristics of Colorectal Signet Ring Cell Carcinoma: A Review. Pathol. Oncol. Res. 27, 1609859 (2021).

12. Fearon, E. R. & Vogelstein, B. A genetic model for colorectal tumorigenesis. Cell 61, 759–767 (1990).

13. Boutin, A. T. et al. Oncogenic Kras drives invasion and maintains metastases in colorectal cancer. Genes Dev. 31, 370–382 (2017).

14. Fernández-Medarde, A. & Santos, E. Ras in cancer and developmental diseases. Genes Cancer 2, 344–358 (2011).

15. Nalli, M., Puxeddu, M., La Regina, G., Gianni, S. & Silvestri, R. Emerging therapeutic agents for colorectal cancer. Molecules 26, 7463 (2021).

16. Infante, J. R. et al. Safety, pharmacokinetic, pharmacodynamic, and efficacy data for the oral MEK inhibitor trametinib: A phase 1 dose-escalation trial. Lancet Oncol. 13, 773–781 (2012).

17. Bao, M. H. R. & Wong, C. C. L. Hypoxia, metabolic reprogramming, and drug resistance in liver cancer. Cells 10, 1715 (2021).

18. Hirpara, J. et al. Metabolic reprogramming of oncogene-addicted cancer cells to OXPHOS as a mechanism of drug resistance. Redox Biol. 25, 101076 (2019).

19. Shanmugasundaram, K. et al. NOX4 functions as a mitochondrial energetic sensor coupling cancer metabolic reprogramming to drug resistance. Nat. Commun. 8, 997 (2017).

20. Allain, E. P., Rouleau, M., Lévesque, E. & Guillemette, C. Emerging roles for UDP-glucuronosyltransferases in drug resistance and cancer progression. Br. J. Cancer 122, 1277–1287 (2020).

21. Hirabayashi, S., Baranski, T. J. & Cagan, R. L. Transformed drosophila cells evade diet-mediated insulin resistance through wingless signaling. Cell 154, 664–675 (2013).

22. Hirabayashi, S. & Cagan, R. L. Salt-inducible kinases mediate nutrient-sensing to link dietary sugar and tumorigenesis in Drosophila. Elife 4, e08501 (2015).

23. Newton, H. et al. Systemic muscle wasting and coordinated tumour response drive tumourigenesis. Nat. Commun. 11, 4653 (2020).

24. Cartee, G. D. & Wojtaszewski, J. F. P. Role of Akt substrate of 160 kDa in insulin-stimulated and contraction-stimulated glucose transport. Appl. Physiol. Nutr. Metab. 32, 557–566 (2007).

25. Na, J. et al. A Drosophila Model of High Sugar Diet-Induced Cardiomyopathy. PLoS Genet. 9, e1003175 (2013).

26. Shimizu, T. et al. The clinical effect of the dual-targeting strategy involving PI3K/AKT/mTOR and RAS/MEK/ERK pathways in patients with advanced cancer. Clin. Cancer Res. 18, 2316–2325 (2012).

27. Tolcher, A. W. et al. Phase I study of the MEK inhibitor trametinib in combination with the AKT inhibitor afuresertib in patients with solid tumors and multiple myeloma. Cancer Chemother. Pharmacol. 75, 183–189 (2015).

28. Bedard, P. L. et al. A Phase Ib dose-escalation study of the oral pan-PI3K inhibitor buparlisib (BKM120) in combination with the oral MEK1/2 inhibitor trametinib (GSK1120212) in patients with selected advanced solid tumors. Clin. Cancer Res. 21, 730–738 (2015).

29. Grilley-Olson, J. E. et al. A phase Ib dose-escalation study of the MEK inhibitor trametinib in combination with the PI3K/mTOR inhibitor GSK2126458 in patients with advanced solid tumors. Invest. New Drugs 34, 740–749 (2016).

30. Shapiro, G. I. et al. Phase Ib study of the MEK inhibitor cobimetinib (GDC-0973) in combination with the PI3K inhibitor pictilisib (GDC-0941) in patients with advanced solid tumors. Invest. New Drugs 38, 419–432 (2020).

31. Ho, M. Y. K. et al. Trametinib, a first-in-class oral MEK inhibitor mass balance study with limited enrollment of two male subjects with advanced cancers. Xenobiotica 44, 352–368 (2014).

32. Glozak, M. A. & Seto, E. Histone deacetylases and cancer. Oncogene 26, 5420–5432 (2007).

33. Li, Y. & Seto, E. HDACs and HDAC inhibitors in cancer development and therapy. Cold Spring Harb. Perspect. Med. 6, a026831 (2016).

34. Lee, J. H. et al. Development of a histone deacetylase 6 inhibitor and its biological effects. Proc. Natl. Acad. Sci. U. S. A. 110, 15704–15709 (2013).

35. Croisy, A., Friesen, M. & Bartsch, H. Species-specific activation of phenacetin into bacterial mutagens by hamster liver enzymes and identification of n-hydroxyphenacetin o-glucuronide as a promutagen in the urine. Cancer Res. 42, 3201– 3208 (1982).

36. Cummings, J., Ethell, B. T., Jardine, L. & Burchell, B. Glucuronidation of SN-38 and NU/ICRF 505 in human colon cancer and adjacent normal colon. Anticancer Res. 26, 2189–2196 (2006).

37. Warburg, O. On the origin of cancer cells. Science (80-.). 123, 309–314 (1956).

38. Wahdan-Alaswad, R. et al. Glucose promotes breast cancer aggression and reduces metformin efficacy. Cell Cycle 12, 3759–3769 (2013).

39. Marcucci, F. & Rumio, C. Glycolysis-induced drug resistance in tumors—A response to danger signals? Neoplasia (United States) 23, 234–245 (2021).

40. Sato, T. & Clevers, H. Primary Mouse Small Intestinal Epithelial Cell Cultures. In Methods in Molecular Biology vol. 945 319–328 (2013).

